# Actin nucleators safeguard replication forks by limiting nascent strand degradation

**DOI:** 10.1101/2023.01.12.523639

**Authors:** Jadwiga Nieminuszczy, Peter R. Martin, Ronan Broderick, Joanna Krwawicz, Alexandra Kanellou, Camelia Mocanu, Vicky Bousgouni, Charlotte Smith, Kuo-Kuang Wen, Beth L. Woodward, Chris Bakal, Fiona Shackley, Andres Aguilera, Grant S. Stewart, Yatin M. Vyas, Wojciech Niedzwiedz

**Affiliations:** Cancer Biology, The Institute of Cancer Research, London, SW3 6JB, UK; Department of Pediatrics, Division of Pediatric Hematology-Oncology, PennState College of Medicine, PennState Health Children’s Hospital, Hershey, Pennsylvania 17033, USA; Genome Stability and Human Disease Laboratory, Institute of Cancer and Genomic Sciences, University of Birmingham, Birmingham B15 2TT, UK; Paediatric Immunology, Allergy and Infectious Diseases, Sheffield Children’s Hospital NHS Foundation Trust, Sheffield, UK; Centro Andaluz de Biología Molecular y Medicina Regenerativa CABIMER, Universidad de Sevilla-CSIC-Universidad Pablo de Olavide, Seville, Spain

**Keywords:** DNA replication, single-stranded DNA, genome stability, actin, actin nucleation

## Abstract

Accurate genome replication is essential for all life and a key mechanism of disease prevention, underpinned by the ability of cells to respond to replicative stress (RS) and protect replication forks. These responses rely on the formation of Replication Protein A (RPA)-single stranded (ss) DNA complexes, yet this process remains largely uncharacterized. Here we establish that actin nucleation-promoting factors (NPFs) associate with replication forks, promote efficient DNA replication and facilitate association of RPA with ssDNA at sites of RS. Accordingly, their loss leads to deprotection of ssDNA at perturbed forks, impaired ATR activation, global replication defects and fork collapse. Supplying an excess of RPA restores RPA foci formation and fork protection, suggesting a chaperoning role for actin nucleators (ANs) (i.e., Arp2/3, DIAPH1) and NPFs (i.e, WASp, N-WASp) in regulating RPA availability upon RS. We also discover that β-actin interacts with RPA directly *in vitro*, and *in vivo* a hyper-depolymerizing β-actin mutant displays a heightened association with RPA and the same dysfunctional replication phenotypes as loss of ANs/NPFs, which contrasts with the phenotype of a hyper-polymerizing β-actin mutant. Thus, we identify components of actin polymerization pathways that are essential for preventing ectopic nucleolytic degradation of perturbed forks by modulating RPA activity.

## INTRODUCTION

Faithful genome duplication during cell division ensures accurate transmission of genetic information to daughter cells. This process relies on the replication of the entire genetic material during S-phase by thousands of replication forks (RF) emanating from numerous origins of replication. However, replication is constantly challenged by endogenous and exogenous insults that may damage RFs (1–3). To respond to replicative stress and prevent stalled forks from collapsing into DNA double-strand breaks (DSBs), a mutagenic lesion if repaired incorrectly, cells have evolved intricate genome surveillance mechanisms underpinned by the ATR and ATM kinases (2,4,5). Mutations in genes involved in the DNA replication and repair (DDR) pathway lead to human disorders that clinically manifest in developmental abnormalities and cancer, underscoring their importance to human health (1,6,7).

A key step in the replication stress response (RSR) is the formation of RPA-coated single stranded DNA (RPA-ssDNA) complex at sites of RF damage. Replication protein A (RPA) is a heterotrimeric complex (RPA1, RPA2, RPA3) with essential roles in several aspects of DNA metabolism, with conservation from yeast to humans. During DNA replication, RPA protects ssDNA transiently formed at perturbed forks against nucleolytic degradation and serves as a platform for recruitment and regulation of several DDR factors, including ATR (8,9). Interestingly, work in yeast *Saccharomyces cerevisiae* (Sc) and Xenopus showed that cells actively control the ability of RPA to associate with ssDNA, suggesting an importance of this process for DNA-associated transactions and genome stability (10,11). Indeed, dysregulation of RPA expression has been implicated in tumour progression, chemotherapy-resistance, or DNA-damage tolerance (12–14). Accordingly, we have recently identified a novel function for the Wiskott-Aldrich syndrome protein (WASp), a regulator of ARP2/3-mediated actin nucleation, in promoting assembly of RPA-ssDNA complexes in *cis* via its direct interaction with RPA(15). Interestingly however, several other factors involved in actin dynamics, for example N-WASp (nucleation-promoting factor, NPF) or ARP2/3 and Formins (actin nucleators, ANs) have all been shown to participate in a range of responses essential for genome maintenance, including DNA replication, centromere maintenance, or DSB repair (16,17), (18–21) (22) (23). However, these factors do not share the same high level of evolutionary conservation for the RPA1-binding motif as WASp (15), and thus underscores the question of how do these NPFs/ANs participate in regulating genome stability?

Here we find that besides WASp, another member of the NPF family N-WASp as well as ANs (ARP2/3, DIAPH1) collectively referred to hence after as (NPF/ANs) are recruited to RFs and function to facilitate the formation/stability of RPA-ssDNA complexes and safeguard genome duplication. Consequently, their loss triggers defective ATR activation, global RF dysfunction and genome instability. Significantly, we discover that monomeric actin (G-actin) interacts directly with RPA, and cells expressing a hyper-depolymerization β-actin (G13R) mutant phenocopy defects associated with NPF/ANs loss i.e., impaired RPA foci formation, impaired ATR activation, and increased fork collapse. In contrast, cells expressing a hyper-polymerization β-actin (S14C) mutant do not show these defects, likely implying a more direct role of actin polymerisation in RF protection. In conclusion, we propose that the association of NPF/ANs with sites of ongoing replication and/or stalled RFs plays a critical role, either directly or indirectly through actin state changes, in facilitating formation of RPA-ssDNA complexes essential for fork protection.

## MATERIAL and METHODS

### Cell lines and drug treatments

HeLa cells were a generous gift from Dr F. Esashi. U2OS cells and U2OS over expressing RPA (Super RPA) cells were a kind gift of Dr L. Toledo. These cell lines were cultured in Dulbecco’s modified Eagle’s medium (DMEM) supplemented with 10% fetal bovine serum (FBS) and standard antibiotics. Primary dermal fibroblasts were maintained in Dulbecco’s modified Eagle’s medium (DMEM; Life Technologies) supplemented with 20% fetal calf serum (FCS), 5% L-glutamine, and 5% penicillin-streptomycin (Invitrogen) antibiotics. Primary fibroblasts were immortalized with 293FT (Invitrogen)-derived supernatant containing a human telomerase reverse transcriptase (TERT) lentivirus that was generated with the plasmids pLV-hTERT-IRES-hygro (gift from Tobias Meyer; Addgene #85140)(24), psPax2 (gift from Didier Trono; Addgene #12260), and pMD2.G (gift from Didier Trono; Addgene #12259). Selection was performed with hygromycin (Invitrogen) at 70 mg/mL. Fibroblast complementation was carried out using a lentiviral vector (pLVX-IRES-Neo; TakaraBio) that encoded 3xHA-tagged DIAPH1 in combination with the lentiviral packaging plasmids described above. Selection was performed with geneticin (Invitrogen) at 400 mg/mL. Expression of HA-tagged DIAPH1 was validated by Western blotting. All cell lines were routinely tested for mycoplasma. Cells were treated with hydroxyurea (Sigma-Aldrich,) with indicated doses. Where indicated cells were pre-treated with 100 μM ARP2/3i (CK-666, Sigma-Aldrich) for 1 hour prior to further drug treatment or harvesting.

### Plasmids

Plasmids encoding YFP NLS Beta-Actin, YFP NLS Beta-Actin S14C and YFP NLS Beta-Actin G13R were a gift from Primal de Lanerolle (Addgene plasmid # 60613; http://n2t.net/addgene:60613; RRID:Addgene_60613; Addgene plasmid # 60614; http://n2t.net/addgene:60614; RRID:Addgene_60614; Addgene plasmid # 60615; http://n2t.net/addgene:60615; RRID:Addgene_60615 respectively)(25). p11d-tRPA(123) plasmid was a kind gift from David Cortez.

### Immunoblotting

Cell lysis was carried out in urea buffer (9 M urea, 50 mM Tris HCL, pH 7.5, 150 mM β-mercaptoethanol) followed by sonication using a soniprep 150 (MSE) probe sonicator or in RIPA buffer (Sigma-Aldrich), R0278 supplemented with 1× SIGMAFAST protease inhibitors (Sigma-Aldrich), S8830 and 1×PhosStop phosphatase inhibitors (Rosche), PHOSS-RO on ice for 15 min followed by centrifugation at 12,000 rpm for 20 min followed by collection of supernatant containing cell lysate. Cell lysates were prepared in SDS loading buffer (2% SDS, 10% (v/v) glycerol, 2% 2-Mercaptoethanol and 62.5 mM Tris-HCl, pH 6.8) followed by boiling at 95 °C for 10 min. Protein concentrations were determined by spectrophotometry using a Denovix ds-11 fx+ device (Denovix) or by the Bradford assay. Samples were resolved by SDS-PAGE and transferred to PVDF or nitrocellulose followed by blocking in 5% low fat milk in 1× TBS/ 0.1% Tween-20 for 1 h at room temperature. Membranes were washed 3 × 5 min in 1x TBS/ 0.1% Tween-20 and incubated overnight at 4 °C in the indicated primary antibodies in 5% low fat milk in 1× TBS/ 0.1% Tween-20 . Membranes were subsequently washed 3 × 5 min in 1× TBS/ 0.1% Tween-20 and incubated in 5% low fat milk in 1× TBS/ 0.1% Tween-20 containing secondary antibodies for 1 h at room temperature. Membranes were subsequently washed 3 × 5 min in 1x TBS/ 0.1% Tween-20 and developed using Immobilon Western HRP Substrate (Millipore, WBKLS0S00) and imaged using the Azure C300, 600 or 280 instruments (Azure biosystems). Primary antibodies used were: α-Tubulin (Sigma, B-5-1-2; T5168, 1:100,000), MCM2 (Abcam, ab4461, 1:10,000), Lamin-B1 (Santa Cruz, sc-30264, 1:1000), N-WASp (Abcam ab187527, 1:1000), DIAPH1 (Bethyl, A300-077A-T, 1:1000; Santa Cruz, sc-373807, 1:500), CHK1 – phospho S345 (Cell Signalling, 133D3; 2348, 1:1000), CHK1 (Santa Cruz, sc-8408, 1:1000), RPA1 (Abcam ab79398), RPA2 (Abcam ab2175; 1:500), Actin (Sigma, A2066, 1:1000), GFP (Roche, 11 814 460 001, 1:500). Secondary antibodies used were anti-mouse IgG-HRP (Dako, P0447, 1:2000) and anti-rabbit IgG-HRP (Dako, P0448, 1:5000).

### Rapid, Efficient And Practical (REAP) cellular fractionation

To fractionate cytoplasmic and nuclear compartments of the cell we carried out REAP fractionation as described previously (26). HeLa cells grown to 70-90% confluence in a 10 cm tissue-culture dish were washed 2 times in ice cold 1x PBS. Then, 1 ml ice cold PBS was added to a 10 cm tissue culture dish and cells were scraped and collected in 1.5 ml microcentrifuge tubes. Microcentrifuge tubes were pulse spun for 10 s and the supernatant was discarded. Cell pellets were then pipetted up and down 5 times in 1 ml ice cold 0.1 % NP-40/ PBS and pulse spun for 10 s in a microcentrifuge. The supernatant was then transferred to a new microcentrifuge tube (Cytoplasmic fraction). The cell pellet was then resuspended by pipetting up and down 5 times in 1 ml ice cold 0.1 % NP-40/ PBS and pulse spun and the supernatant was discarded. The nuclear pellet was then resuspended in 50 μl of PBS and sonicated for 30 s at 10% amplitude using a soniprep 150 (MSE) probe sonicator. The protein concentration of the cytoplasmic and nuclear fractions were then quantified by Bradford assay and processed by SDS-PAGE and western blot transfer in advance of immunoblot analysis.

### YFP-NLS-β-Actin mammalian expression construct transfection

For YFP-NLS-β-Actin mammalian expression construct transfection, 200,000 cells were seeded into a well of a 6-well tissue culture plate. Cells were incubated at 37°C for 24 hours to allow them to adhere. Per condition, 0.75μg of YFP-NLS-β-Actin mammalian expression construct was transfected following the Lipofectamine 3000 manufacturer’s protocol. Cells were then incubated at 37°C for 24 hours and treated with hydroxyurea at the indicated doses before harvesting cells for down-stream assays.

### Cell Survival Assay

Alamar Blue survival assays were performed in accordance with the manufacturer’s recommendations (Bio Rad). Briefly, 500 cells per well in 96-well plates were plated and untreated or treated with indicated doses of hydroxyurea, cis-platin or Mitomycin C and incubated for 7 days. Alamar blue reagent was added to each well and fluorometric measurements taken after 4h incubation at 37°C.

### RNAi treatment

siRNAs used in this study were as follows: siWASp – GAGUGGCUGAGUUACUUGC and ON-TARGETplus Human WAS siRNA smart pool (LQ-028294-00-0005, Dharmacon); siN-WASp - ON-TARGETplus Human WASL siRNA smart pool (L-006444-00-0005, Dharmacon), CAGCAGAUCGGAACUGUAU (J-006444-07, Dharmacon), UAGAGAGGGUGCUCAGCUA (J-006444-08, Dharmacon); siDIAPH1-ON-TARGETplus Human DIAPH1 siRNA smart pool (L-010347-00-0005, Dharmacon); siRNA targeting luciferase - CGTACGCGGAATACTTCGA (27) was used as control siRNA. Oligonucleotides were transfected using HiPerfect reagent (Qiagen) at 25nM concentration, according to the manufacturer’s protocol. First pulse of siRNA was followed with 2^nd^ pulse after 24 hours and all experiments were performed at maximum knock down-efficiency 72 hours post 1^st^ siRNA pulse.

### Immunofluorescence microscopy

Cells were grown and treated on circular 13 mm diameter coverslips, thickness 1.5 mm. For visualization of 53BP1 foci in all cells and in Cyclin A-positive cells, cells were fixed with 4% paraformaldehyde in PBS for 10 min at room temperature, washed twice in PBS and permeabilised with 0.2% Triton X-100 in PBS for 10 min at room temperature. Coverslips were washed 3 x in PBS cells and were blocked in 10% FBS in PBS for 1 h at room temperature before incubation with primary antibodies in 0.1% FBS in PBS overnight at 4 °C. Coverslips were then washed 4 × 5 min in PBS followed by incubation with secondary antibodies for 1 h at room temperature. Slides were then washed 4 × 5 min in PBS and subsequently mounted with Vectashield mounting medium (Vector Laboratories) with DAPI. Micronuclei were quantified by assessing DAPI stained nuclei. For visualization of RPA2 foci and BrdU foci cells were pre-extracted on ice for 2 min in CSK buffer (10 mM PIPES pH6.8, 300 mM sucrose, 100 mM NaCl, 1.5 mM MgCl_2_, 0.5% TritonX-100), washed with PBS and fixed with 4 % paraformaldehyde in PBS for 10 min at room temperature. Coverslips were then blocked and processed as above. Primary antibodies employed for immunofluorescence were as follows: 53BP1 (MAB3802, Millipore, 1:1000), Cyclin A (ab19150, Abcam, 1:500), RPA2 (NA-18, Calbiochem, 1:200), RAD51 (ab133534, Abcam 1:500), BrdU (BU-1; RPN202 GE Healthcare Life Sciences 1:200). Secondary antibodies for immunofluorescence were as follows: Alexa Fluor 488 anti-rabbit (A21206, Invitrogen, 1:200) and Alexa Fluor 555 anti-mouse (A31570, Invitrogen, 1:400). Images were acquired using a Leica SP8 confocal microscope with a 63× oil objective, using a Zeiss Axio Observer Z1 Marianas™ Microscope attached with a CSU-W spinning disk unit using either a Hamamatsu Flash 4 CMOS camera or a Photometrics Prime 95b sCMOS camera built by Intelligent Imaging Innovations (3i) or a Zeiss Axio Observer Z1 Marianas™ Microscope attached with a CSUX1 spinning disk unit and Hamamatsu Flash 4 CMOS camera built by Intelligent Imaging Innovations (3i). Quantification was carried out using FIJI (ImageJ) software and CellProfiler (Broad Institute).

### iPOND

iPOND was performed as decribed previously (28). Briefly, logarithmically growing HeLa S3 cells (1 × 10^6^ per ml) or HEK293TN cells were incubated with 10 μM EdU for 10 or 15 minutes respectively. Following EdU labeling, cells were fixed in 1% formaldehyde, quenched by adding glycine to a final concentration of 0.125 M and washed three times in PBS. Collected cell pellets were frozen at −80°C and cells were permeabilized by resuspending 1.0–1.5 × 10^7^ cells per ml in ice cold 0.25% Triton X-100 in PBS and incubating for 30 minutes. Before the Click reaction, samples were washed once in PBS containing 0.5% BSA and once in PBS. Cells were incubated for 2 hours at room temperature in Click reaction buffer containing 10 μM azide-PEG(3+3)-S-S-biotin conjugate (Click ChemistryTools, cat. no AZ112-25), 10 mM sodium ascorbate, and 1.6 mM copper (II) sulfate (CuSO4) in PBS. The ‘no Click’ reaction contained DMSO instead of biotin-azide. Following the Click reaction, cells were washed once in PBS containing 0.5% BSA and once in PBS. Cells were resuspended in lysis buffer (50 mM Tris–HCl pH 8.0, 1% SDS) containing protease inhibitor cocktail (Sigma) and sonicated. Samples were centrifuged at 14,500 rcf. at 4°C for 30 minutes and the supernatant was diluted 1:3 with TNT buffer (50 mM Tris pH 7.5, 200 mM NaCl and 0.3% Triton X-100) containing protease inhibitors. An aliquot was taken as an input sample. Streptavidin–agarose beads (Novagen) were washed three times in TNT buffer containing protease inhibitor cocktail. Two hundred microliters of bead slurry was used per 1× 108 cells. The streptavidin–agarose beads were resuspended 1:1 in TNT buffer containing protease inhibitors and added to the samples, which were then incubated at 4°C for 16 hours in the dark. Following binding, the beads were then washed two times with 1 mL TNT buffer, two times with TNT buffer containing 1M NaCl, two times with TNT buffer and protein–DNA complexes were eluted by incubating with 5 mM DTT in TNT buffer. Cross-links were reversed by incubating samples in SDS sample buffer at 95°C for 20 minutes. Proteins were resolved on SDS-PAGE and detected by immunoblotting using specific antibodies.

iPOND in ND1 cells was performed as described(29) with some modifications. Briefly, T cells (~80 million cells per sample) were incubated with 20 μM EdU in 30 ml culture medium at 37°C for 20 min and then washed with fresh medium at 37°C. For HU or thymidine pluse, cells were resuspended either with 30 ml fresh medium at 37 °C as the control or 4 mM HU or 20 μM thymidine in fresh medium. These cells were further cultured at 37°C for another 2 hrs. After labeling/treatments, each cell samples were washed with PBS, and crossed-linked in 1% Formaldehyde in PBS for 20 min at RT, quenched with glycine at the final concentration of 0.125 M for another 5 min, and washed 2 times in PBS. Cell pellets were permeabilized with 10 ml permeabilizing buffer (0.3% Triton-X/0.5% BSA in PBS) 30 min at RT and washed with 0.5% BSA/PBS. Each cell pellet was resuspended in 10 ml PBS as the control, or 10 ml fresh prepared Click buffer (10 mM Sodium ascorbate, 2 mM CuSO4, and 20 μM Biotin-dPEG7-azide) and incubated for 1-2 hr at room temperature (RT). Cells were washed with 0.5% BSA/PBS and then pellets either frozen at −80°C or immediately used for lysis. Each cell pellet was resuspended in 0.8 ml lysis buffer (25 mM NaCl, 2 mM EDTA, 50 mM Tris pH 8, 1% IGEPAL CA630, 0.2 % SDS, 0.5% sodium deoxycholate, and 1X Halt protease and phosphatase inhibitor cocktail (Thermo Fisher)) and incubated for 10 min on ice. Samples were sonicated with a Branson 250 using the settings, 20-25 W, 20 sec constant pulse, and 40 sec pauses for a total of 4 min on ice. Cell lysates were centrifuged at 18k X g for 10 min at RT. The supernatants were collected and diluted with the dilution buffer (the lysis buffer without SDS or sodium deoxycholate). Streptavidin-agarose beads (Millipore Sigma) (80μl/sample) were washed with the dilution buffer 3 times, and then incubated with the diluted samples overnight at 4°C. The beads were again washed 3 times with RIPA buffer. Captured proteins were separated from beads by incubating beads in 50 μl 2× Laemmli Sample Buffer (Bio-Rad) at 95°C for 25 min. The supernatant was collected, and proteins resolved on 4-15% SDS-PAGE and detected by immunoblotting.

### EdU labeling of nascent DNA and Proximity Ligation Assay

Analysis of the association of proteins to nascent DNA by EdU labelling and the Proximity Ligation Assay was carried out as previously described (28,30). Briefly, cells grown on coverslips were labeled with 10 mM EdU for 10 min followed by treatment with 1mM or 4mM hydroxyurea at various timepoints as indicated. Fixation was carried out with 3% formaldehyde, 2% sucrose in PBS for 10 min at room temperature. Slides were then washed twice with PBS and incubated with blocking solution (3% BSA in PBS) for 30 min. Following this, slides were washed 2× in PBS before EdU-Biotin Azide conjugation by click chemistry using the Click-iT reaction (Click-iT assay kit (Thermo Fisher, according to the manufacturer’s instructions). Coverslips were washed 2× with PBS before primary antibody incubation overnight at 4C in 1% BSA/0.1% saponin in PBS. Following primary antibody incubation coverslips were washed 2× in PBS and then the proximity ligation assay was carried out (Duolink *In Situ* Red Starter kit (Sigma Aldrich) according to the manufacturer’s instructions). Coverslips were mounted using Vectashield containing DAPI. Antibodies employed for the PLA assay were as follows: Biotin (Bethyl Laboratories, A150-109A, 1:3000), (Biotin (Jackson Immunoresearch, 200-002-211, 1:1000), WASp (Santa Cruz, sc-5300, 1:500), N-WASp (Abcam, ab187527, 1:500), PCNA (Santa Cruz, sc-56, 1:500) and PCNA (Abcam, ab18197, 1:500). Images were acquired using a Zeiss Axio Observer Z1 Marianas™ Microscope attached with a CSU-W spinning disk unit using either a Hamamatsu Flash 4 CMOS camera or a Photometrics Prime 95b sCMOS with a 63× objective. Image analysis was carried out with FIJI (ImageJ) software.

### DNA fibre analysis

DNA fibre assay was performed as described previously with some modifications (28,31,32). In brief, exponentially growing cells were first incubated with 25 μM iododeoxyuridine (IdU) and then with 125 μM chlorodeoxyuridine (CldU) for the indicated times. Fibre spreads were prepared from 0.5 x10^6^ cells/ml. Slides were stained as described previously (31,32). Images were acquired with Leica SP8 or Carl Zeiss LSM 710 Meta confocal microscope using a 63× oil objective. Analysis was performed using the ImageJ software package (National Institutes of Health). A minimum of 100 fibres from three, unless stated otherwise, independent experiments were scored. Mann-Whitney test was used to determine statistical significance.

### QPCR

RNA extraction and cDNA synthesis were performed using the RNeasy mini kit (Qiagen) and High Capacity cDNA Reverse Transcription Kit (Applied Biosystems). QPCR analysis was performed with QuantStudio™ 6 Flex Real-Time PCR System (Applied Biosystems) using SYBR Green PCR Master Mix (Life Technologies) and the following primers GTCCTACTTCATCCGCCTTTAC and TCGTCTGCAAAGTTCAGCC for WASP and GGCATGGACTGTGGTCATGAG and TGCACCACCAACTGCTTAGC for GAPDH.

### RPA complex expression, purification and in vitro pull down

RPA complex was expressed from a plasmid (a kind gift from David Cortez (Vanderbilt University, Nashville, Tennessee, US) (33,34) encoding 6xHis-tagged RPA1 (70 kDa), tagless RPA2 (32 kDa), and 6×His-tagged RPA3 (14 kDa) in BL21 (D3) *E. coli* cells for 2 h at 37°C in LB medium with 100 μM/ml ampicillin after induction with 1 mM IPTG. Cells were harvested and lysed in buffer containing 50 mM Tris-HCl, pH 8.0, 400 mM NaCl, 1 mM PMSF, 10 μM ZnCl_2_, 5% glycerol and protease inhibitor cocktail (Sigma, #S8830). Supernatant was applied to a HisTrap HP 5 ml column (DE Healthcare) calibrated in 5 volumes of calibration buffer containing 20 mM Tris-HCl, pH 8.0, 500 mM NaCl, 1 mM PMSF, 10 μM ZnCl_2_ and 5% glycerol. Next, the column was washed in 10 column volumes with washing buffer: 20 mM Tris-HCl, pH 7.6, 500 mM NaCl, 20 mM imidazole, 10 μM ZnCl2 and 5% glycerol. Then, the proteins were eluted from column with the linear imidazole gradient in final concertation as follow, 20 mM Tris-HCl, pH 7.5, 100 mM NaCl, 300 mM imidazole, 10 μM ZnCl_2_ and 5% glycerol. The RPA complex of three proteins was bound to the column and was eluted from it by high imidazole concentration (about 250 mM). The eluate was diluted to bring the NaCl concentration down to 50 mM. It was then applied to a HiTrap Heparin HP 5 ml column (GE Healthcare) in 20 mM Tris-HCl, pH 7.5, 50 mM NaCl, 1 mM PMSF, 10 μM ZnCl_2_ and 5% glycerol, and eluted with 10 volumes of 50–1000 mM NaCl linear gradient. The fractions containing all three proteins were joined together and purified RPA complex was rebuffered into storage buffer (50 mM Tris-HCl, pH 8.0, 100 mM NaCl, 1 mM DTT, 5% glycerol), aliquoted and frozen in −80°C. The complex composition was confirmed by western blot using antibodies recognizing His-tag peptide and RPA2 protein and by binding RPA complex to ssDNA and lack of biding it to dsDNA. Interaction of RPA complex with monomeric actin was analysed using G-Actin Sepharose beads (Hypermol). Reaction was performed in a buffer containing (10mM Tris-HCl pH7.5, 150 mM NaCl, 1 mM DTT, 5% glycerol, 0.1 % NP-40). Complexes were extensively washed with IP buffer before elution in 2X SDS sample buffer and subsequently boiled for 3 min followed by centrifugation. The resultant supernatant fraction was retained as the eluate.

### YFP-NLS-β-Actin Co-immunoprecipitations

YFP-NLS-β-Actin expression constructs were transfected into HEK293TN cells using lipofectamine 2000 (Thermo Scientific) following manufacturer’s instructions. HEK293TN cells were harvested 48 hours post transfection from a 70%-80% confluent 15 cm tissue culture dish by trypsinisation, inactivation with serum containing media, centrifugation at 1500 rpm for 5 min, followed by the removal of the supernatant. Cells were washed twice in phosphate-buffered saline and lysed in IP buffer 1 (100 mM NaCl, 0.2% Igepal CA-630, 1 mM MgCl_2_, 10% glycerol, 5 mM NaF, 50 mM Tris-HCl, pH 7.5), supplemented with SigmaFAST Protease Inhibitor Cocktail, EDTA free (Sigma-Aldrich, S8830) and 25 U ml–1 Benzonase (Novagen). After nuclease digestion, NaCl and EDTA concentrations were adjusted to 200 mM and 2 mM, respectively, and lysates were cleared by centrifugation (16,000 × g for 25 min). Lysates were incubated for 1 hr with 20 μl of agarose binding control beads (ChromoTek), equilibrated in IP buffer 2 (200 mM NaCl, 0.2% Igepal CA-630, 1 mM MgCl_2_, 10% glycerol, 5 mM NaF, 2 mM EDTA 50 mM Tris-HCl, pH 7.5) to pre-clear. The agarose binding control beads were pelleted by centrifugation at 2000 *xg* for 2 min and total protein quantified by the Bradford assay. Lysates were then incubated with 20 μl of GFP-Trap agarose beads (ChromoTek), equilibrated in IP buffer 2, for 2 hours with end-to-end mixing at 4 °C at a concentration of 1mg/ml. Complexes were washed 4× in IP buffer 2 before resuspension in 2× SDS sample buffer and elution by boiling at 95°C for 10 min.

### STRING analysis

Gene accession codes from IPOND analysis were input into the multiple protein search function of STRING (https://string-db.org/cgi/input?sessionId=b5U7THWxoyjd&input_page_active_form=multiple_identifiers) with the detected organisms selected as Homo sapiens. The network was then exported from STRING and imported into Cytoscape (v3.8.2). Functional enrichment network analysis was then conducted using the STRING plug-in, with the network to be set as background set to genome. The STRING Enrichment tab was selected, and the generated network was filtered by GO Biological processes. GO Biological processes associated with all the actin related processes were selected and exported as a new network. The subsequently generated network was saved, and the file was uploaded to STRING (https://string-db.org/cgi/input?sessionId=b5U7THWxoyjd&input_page_show_search=on). Active interaction sources settings were adjusted to display links associated with only experiments and databases. The minimum required interaction score was adjusted to high confidence (0.700) and the network was updated. GO terms GO:0034314 Arp2/3 complex-mediated actin nucleation and GO:0008154 Actin polymerization or depolymerization were selected in the analysis tab functional enrichment biological process list.

### Statistics and reproducibility

Statistical analyses were done using GraphPad Prism 9 (GraphPad Software Inc.) or Microsoft Excel. Unpaired Students’ t-test or Mann Whitney test were used to determine statistical significance as indicated in the Figure Legends.

### Contact for reagent and resource sharing

Further information and requests for resources and reagents should be directed to the Lead Contact, Wojciech Niedzwiedz (wojciech.niedzwiedz.icr.ac.uk)

## RESULTS

### WASp, N-WASp, and DIAPH1 are recruited to sites of DNA replication and protect cells from replicative stress

To address the possible role of NPF/ANs during normal and stressed DNA replication we performed a deep mining of our iPOND (isolation of proteins on nascent DNA) data set (28,35,36) combined with STRING analysis (http://string-db.org) performed with or without hydroxyurea (HU)-mediated replicative stress (RS) in HeLa cells. This analysis identified a cluster of proteins involved in actin polymerisation, including the ARP2/3 complex, WASp family proteins, and members of the formin family of actin nucleators (DIAPH1, 2 and 3), but not Spire family proteins, the latter known to nucleate actin polymers directly without Arp2/3 (Figure 1A). We validated the presence of these factors at replication forks (RFs) in two ways: a) iPOND/Western blot (WB) analysis and b) EdU-based proximity ligation assay (PLA), aka quantitative *in situ* analysis of protein interaction at RFs (SIRF) ^15 27, 30^. Using iPOND/WB analysis we could readily detect WASp, N-WASp, and DIAPH1, as well as RPA2 (control) enriched at HU-perturbed RFs in both human T cells and HEK293TN cells (Supplementary Figures S1A-C). Notably, a thymidine chase experiment showed increased association with perturbed forks, which indicates that they are likely to be recruited to RFs in a RSR dependent manner (29,37) (Supplementary Figures S1A-C). Accordingly, utilizing PLA we could also readily detect proximity of the NPFs (N-WASp, WASp) to newly synthesised DNA (monitored by EdU labelling of nascent DNA marking RFs) in HeLa cells. The nuclear PLA signal for N-WASp and WASp was significantly enriched upon HU-induced RS compared to untreated as well as control samples (Figure 1B). To ascertain that the observed phenotype is not restricted to EdU labelling we repeated these analyses using an antibody against PCNA - an established marker of RFs. Again, we readily detected association between PCNA and N-WASp or WASp (Supplementary Figure S1D and E) confirming their proximity to sites of DNA replication, which is in line with our recent analysis demonstrating WASp localisation to RFs in human T and B cells (15). Moreover, in support of our iPOND data, thymidine addition resulted in a significant decrease in PLA signal between N-WASp or WASp and nascent DNA (Figure 1C). Taken together, these data suggest that members of the NPF/AN family of proteins enrich at perturbed RFs in human immune and nonimmune cells, malignant or non-malignant.

**Figure 1.**
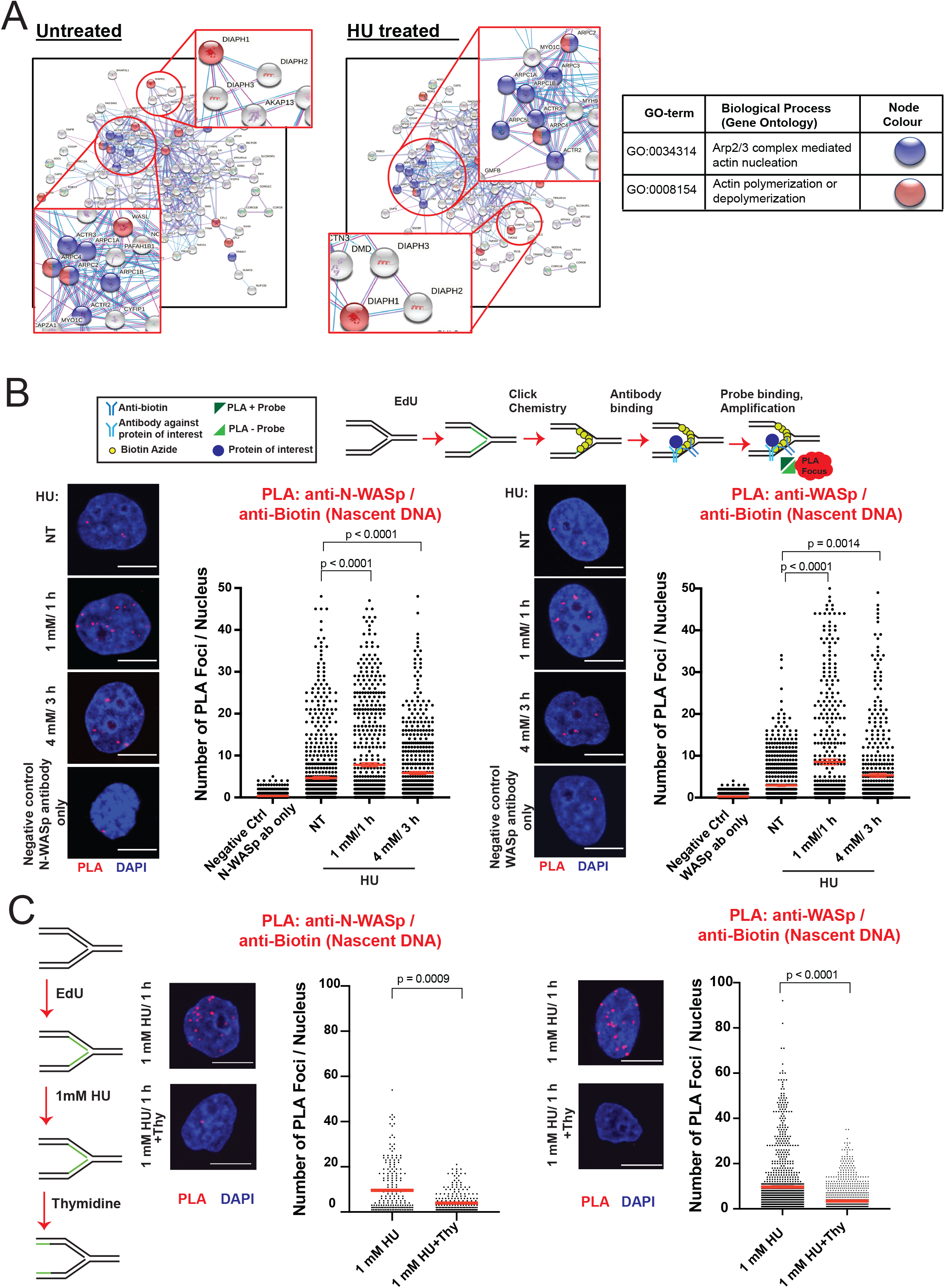
WASp, N-WASp and DIAPH1 associate with replication forks. A. STRING analysis of iPOND/MS data in untreated and HU treated HeLa S3 cells. B. Representative images and dot plots of the number of N-WASp/Biotin (Nascent DNA) and WASp/Biotin (Nascent DNA) PLA foci per nucleus in HeLa cells upon HU treatment as indicated (red lines indicate mean values). Dot plots represent data pooled from three independent experiments. Statistical significance was determined using the Mann-Whitney test. C. Representative images and dot plots of number of N-WASp/Biotin (Nascent DNA) and WASp/Biotin (Nascent DNA) PLA foci per nucleus in HeLa cells treated with 1mM HU for 1h, followed or not by thymidine chase as indicated (red lines indicate mean values). Dot plots represent data pooled from two independent experiments. Statistical significance was determined using the Mann-Whitney test.

### Deficiency in NPF/ANs provokes global replication dysfunction

To address the mechanism by which NPF/ANs manage RS, we employed DNA fibre assay to analyse various aspects of stressed DNA replication (38,39). Strikingly, replicative stress induced by HU, resulted in a significant reduction in the average fork velocity as well as impaired fork restart in cells transiently depleted of N-WASp, DIAPH1 or WASp by RNA interference relative to WT control (Figure 2A, Supplementary Figures S2A-C). To validate the specificity of siRNA data we recapitulated these observations using (i) multiple different siRNAs against N-WASp or WASp (Supplementary Figure S2D), (ii) patient-derived DIAPH1-deficient cell-line and a complemented control cell line (Figure 2B), as well as (iii) a specific ARP2/3 inhibitor (CK-666; Supplementary Figure S2E). In addition, loss of WASp, N-WASp or DIAPH1 significantly impaired fork restart upon HU treatment (Figure 2C). WASp family members function as positive regulators of ARP2/3-mediated generation of short branched F-actin filaments (40) whereas formins, of the DIAPH-family, nucleate long unbranched F-actin filaments (41). Thus, given the striking similarity of the replication phenotypes we observed in the tested NPF/ANs, we next tested if these factors function in a single or parallel pathway(s) to manage RS. To this end, we tested the impact on DNA replication (measured as fork speed) of either WASp or DIAPH1 depletion alone or together with an ARP2/3 inhibitor under HU induced stress. Targeting either WASp or ARP2/3 alone as well as inhibiting both together resulted in a similar replication defect, indicating that WASp and ARP2/3 both function in the same pathway to manage RS (Supplementary Figure S3A). In contrast, inhibition of ARP2/3 in DIAPH1-deficient cells led to a significantly greater replication defect than the defect displayed by DIAPH1-deficient cells alone (Figure 2D), indicating an additive effect between branched and unbranched actin polymerisation pathways in promoting DNA replication. Taken together, these data suggest that WASp:ARP2/3 and DIAPH:formin mediated actin signalling is required to facilitate fork stability and global RF dynamics.

**Figure 2.**
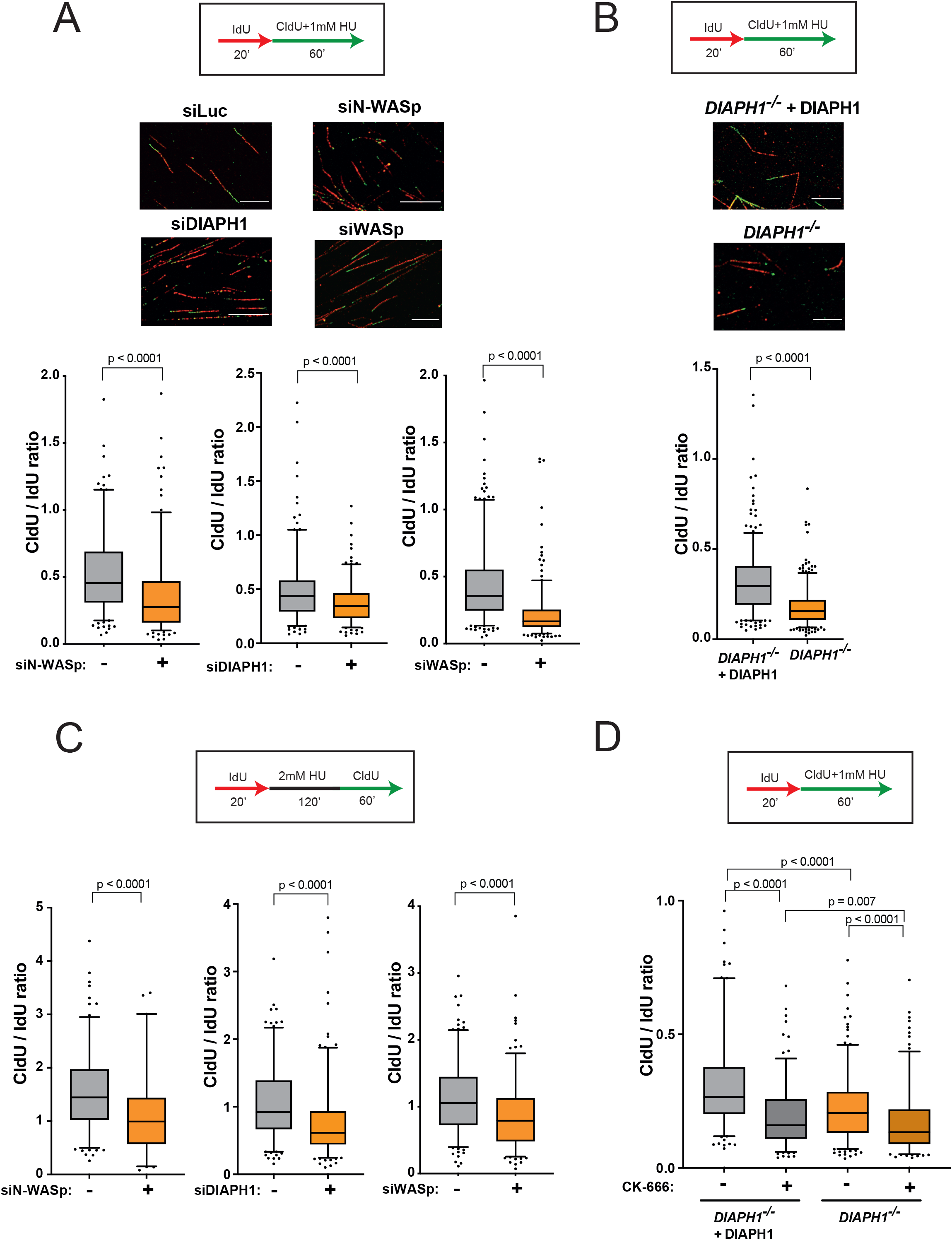
WASp, N-WASp, ARP2/3, and DIAPH1 are required for intact replication dynamics under condition of replicative stress. A. Representative DNA fibre images and box plots of CldU/IdU tract ratios from HeLa cells treated with control siRNA (-) or siRNA against WASp, N-WASp or DIAPH1(+) and exposed to 1mM HU during CldU labelling pulse. Whiskers indicate 5-95 percentile. Box plots represent data pooled from three (siWASp) or two (siN-WASp and siDIAPH1) independent experiments. Statistical significance was determined using the Mann-Whitney test. B. Box plots of CldU/IdU tract ratios from DIAPH1-deficient patient-derived cells treated with 1mM HU during CldU labelling pulse. Whiskers indicate 5-95 percentile. Box plots represent data pooled from three independent experiments. Statistical significance was determined using the Mann-Whitney test. C. Box plots of CldU/IdU tract ratios of HeLa cells treated with control siRNA (-) or siRNA against N-WASp, DIAPH1 or WASp (+). After IdU pulse, cells were exposed to 2mM HU for 2h followed by release into CldU for 60 minutes. Whiskers indicate 5-95 percentile. Box plots represent data pooled from two independent experiments. Statistical significance was determined using the Mann-Whitney test. D. Box plots of CldU/IdU tract ratios from DIAPH1-deficient patient-derived cells treated with CK-666 (ARP2/3 inhibitor) (+) or DMSO (-). During CldU labelling pulse, cells were exposed to 1mM HU. Whiskers indicate 5-95 percentile. Box plots represent data pooled from two independent experiments. Statistical significance was determined using the Mann-Whitney test.

### Interfering with actin polymerisation elicits replication-associated DNA damage

Unstable RFs are prone to collapse into single-ended DSBs. Thus, to determine the fate of HU-stalled RFs when actin nucleation is impaired, we analysed the formation of 53BP1 foci (a canonical marker of DSB) in cells depleted for either N-WASp, WASp or DIAPH1, either in unperturbed conditions, after 1 mM/ 1h HU treatment (mild RS associated with fork stalling) or 4 mM/ 3h HU treatment (high RS, associated with elevated RF collapse)(42). We noticed a significant increase in the number of 53BP1 foci in cyclin A-positive (S/G2-phase) cells after depleting either of the three proteins as compared to their respective controls, most likely due to extensive fork collapse (Figure 3A). Notably, we reproduced these findings in cells treated with an ARP2/3 inhibitor (CK-666) and in the patient-derived DIAPH1-deficient cells (Supplementary Figure S3B). Fork collapse is known to induce genome instability resulting in the increased formation of micronuclei, which are particularly prominent in cancer cell models (e.g., HeLa) that display an intrinsically high levels of replicative stress. Consistently, loss of WASp, N-WASp, or DIAPH1 as well as inhibition of ARP2/3 resulted in increased levels of micronuclei compared to control cells even in the absence of any exogenous replication stressors (Figure 3B and Supplementary Figure S3C). This observation further underscores the role of NPF/ANs in preventing genome-instability not only in response to exogenous genotoxic insults but also those arising from endogenous sources of RS, such as R loops (RNA-DNA hybrids), transcription-replication (T-R) conflicts, etc (15). Collectively, these data support the notion that NPF/ANs promote recovery of perturbed RFs and suppress replication-associated genome instability during normal DNA replication as well as in response to replicative stress.

**Figure 3.**
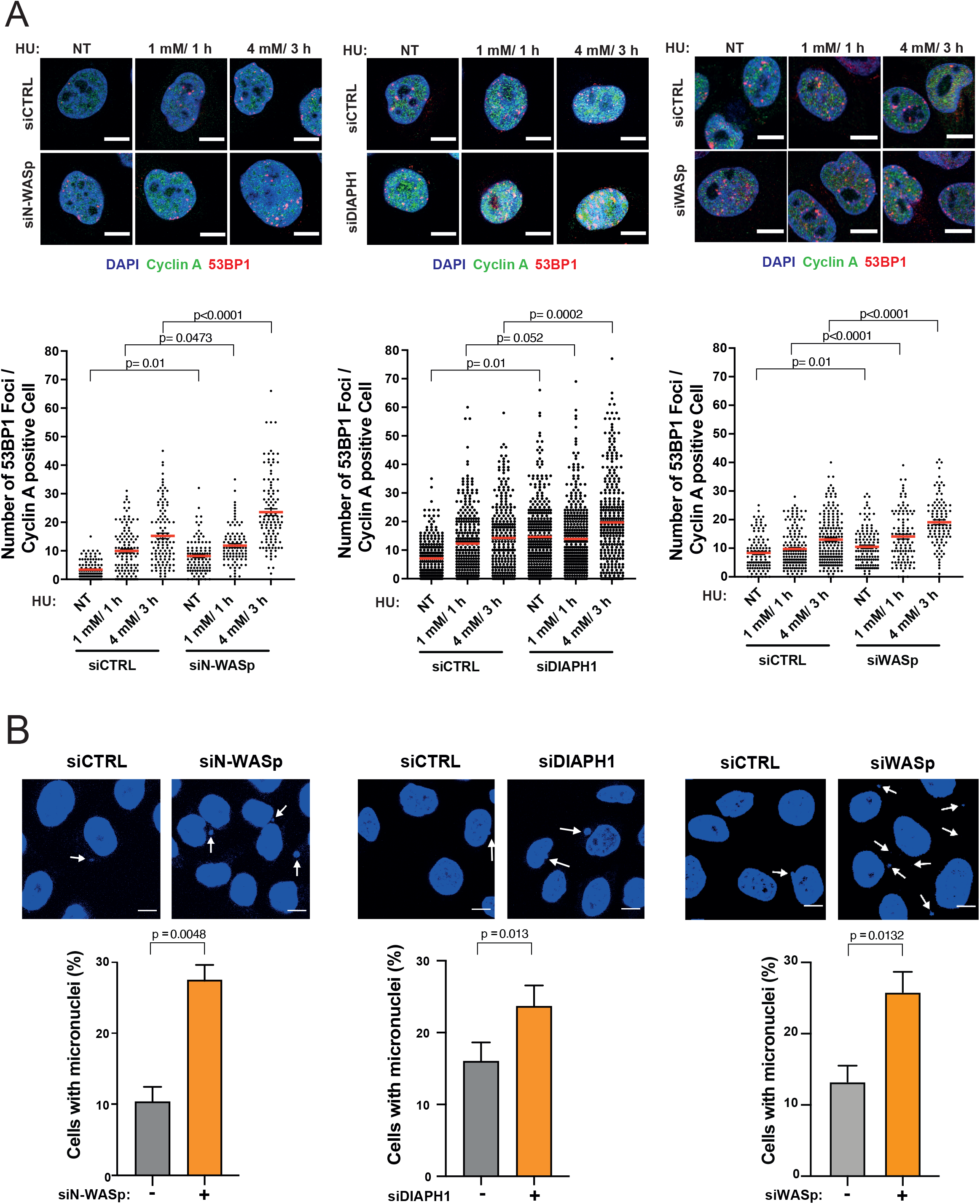
Depletion of WASp, N-WASp or DIAPH1 leads to increase in replication fork collapse. A. Representative images and dot plots of the number of 53BP1 foci per cyclin A positive HeLa cells treated with control siRNA (siCTRL) or siRNA targeting N-WASp, DIAPH1 or WASp upon HU treatment. Dot plots represent data pooled from three independent experiments. Statistical significance was determined using the Mann-Whitney test. Representative images and quantification of micronuclei formation in HeLa cells treated with control siRNA (-) or siRNA targeting N-WASp, DIAPH1 or WASp (+). Dot plots represent data pooled from at least three independent experiments. Bar charts represent mean +/− SEM of three experiments and statistical significance was determined using t test.

### Actin nucleation protects replication forks from uncontrolled degradation by promoting RPA localisation to ssDNA

Since unstable RFs undergo excessive nucleolytic degradation (43–46), we tested whether loss of NPF/ANs influences this process by employing a modified DNA fibre protocol (28). Interestingly, depletion of N-WASp, DIAPH1 or WASp in HeLa cells, resulted in a significant shortening of the CldU tracts (in the IdU>CIdU>HU labelling scheme) compared to controls (Figure 4A). This data indicate that pathological nucleolytic degradation of HU-perturbed forks is not restricted to WASp deficiency or to human T and B lymphocytes, but is a more general phenotype associated with depletion of multiple NPF/AN proteins(15). As pathological fork resection is attributed to the unrestrained activity of the MRE11 nuclease, we monitored fork degradation in cells treated with the MRE11 inhibitor mirin. Treatment with mirin supressed excessive fork degradation observed in cells depleted of NPF/ANs (Figure S3D). Moreover, depletion of SMARCAL1 also rescued excessive fork degradation in cells depleted of N-WASp, WASp or DIAPH1 suggesting a role in fork protection downstream to RF reversal (Figure 4B). Since pathological fork resection is attributed to fork deprotection (47) and we have shown that WASp-deficient immune cells (T and B cells) display defective ssDNA-RPA complex formation(15) we tested if depletion of other NPF/ANs would have a similar function in non-immune cells (i.e. HeLa) as well. Remarkably, depletion of either N-WASp or DIAPH1 or WASp alone led to defective RPA foci formation upon HU-induced RS (Figure 4C, Supplementary Figure S4A). To discount the possibility of off-target effects associated with siRNA, we again validated the specificity of our data by recapitulating these observations in two ways: (i) employing different siRNAs against N-WASp or WASp (Supplementary Figure S4B) as well as (ii) utilising a specific ARP2/3 inhibitor (CK-666; Figure S4C). In line with this, loading of RAD51 at RFs, an event dependent on RPA-ssDNA formation (8) was also defective, as measured by the PLA assay between nascent DNA (EdU labelling) and RAD51 (Figure 5A). Furthermore, the incidence of RAD51 foci detected by IF after HU treatment upon WASp depletion was also significantly reduced when compared to those treated with control siRNA (Figure 5B), thus supporting our PLA-based analysis. Formation of RPA-ssDNA is required for ATR activation and consequently, we have shown recently that WASp is required for this response in T and B cells(15). To ascertain whether other members of the NPF/ANs family are similarly required for this response we analysed the efficiency of ATR signalling in cells depleted for N-WASp or DIAPH1. Again, and in line with the defective RPA loading onto ssDNA, ATR activity was significantly compromised as evidenced by impaired phosphorylation of CHK1 (pSer345) and increased replication origin firing in N-WASp, WASp and DIAPH1 depleted HeLa cells, and also in DIAPH1-deficient cells (Figure 5C and D).

**Figure 4.**
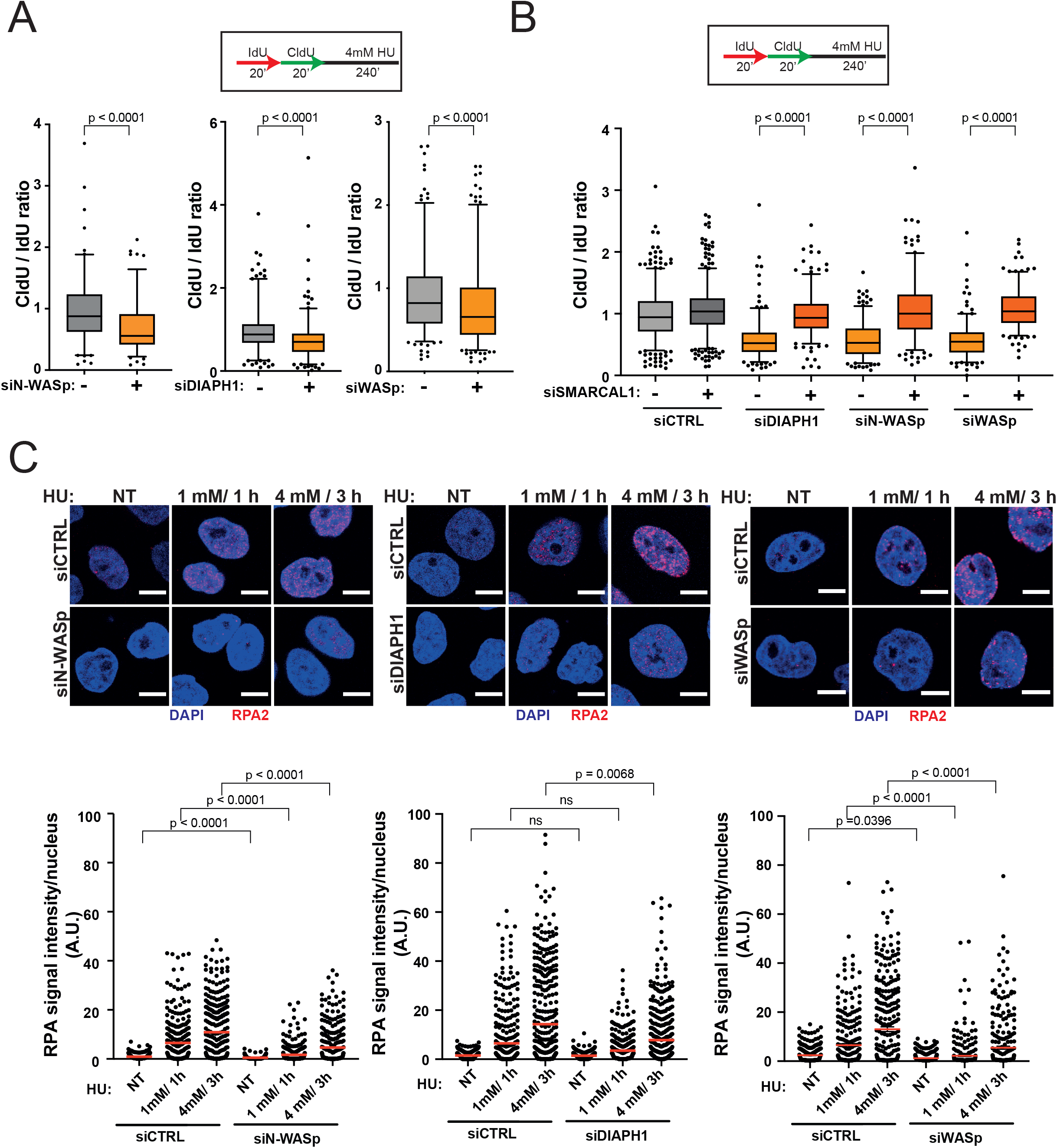
Depletion of WASp, N-WASp or DIAPH1 leads to increased degradation of nascent DNA and defects in RPA recruitment to perturbed replication forks. A. Box plots of CldU/IdU tract ratios from HeLa cells treated with control siRNA(-) or siRNA targeting N-WASp, DIAPH1 or WASp (+). CldU/IdU pulse labelling was followed by treatment with 4mM HU for 4h. Whiskers indicate 5-95 percentile. Box plots represent data pooled from four (siWASp), or two (siN-WASp, siDIAPH1) independent experiments. Statistical significance was determined using the Mann-Whitney test. B. Box plots of CldU/IdU tract ratios from HeLa cells treated with control siRNA or siRNA targeting N-WASp, DIAPH1 or WASp and SMARCAL1. CldU/IdU pulse labelling was followed by treatment with 4mM HU for 4h. Whiskers indicate 5-95 percentile. Box plots represent data pooled from two independent experiments. Statistical significance was determined using the Mann-Whitney test. C. Representative images and dot plots of RPA signal intensity per nucleus of Hela cells treated with control siRNA or siRNA targeting N-WASp, DIAPH1 or DIAPH1 upon HU treatment as indicated (red lines indicate mean values). Dot plots represent data pooled from three (siN-WASp and siWASp) or three (siDIAPH1) independent experiments. Statistical significance was determined using the Mann-Whitney test.

**Figure 5.**
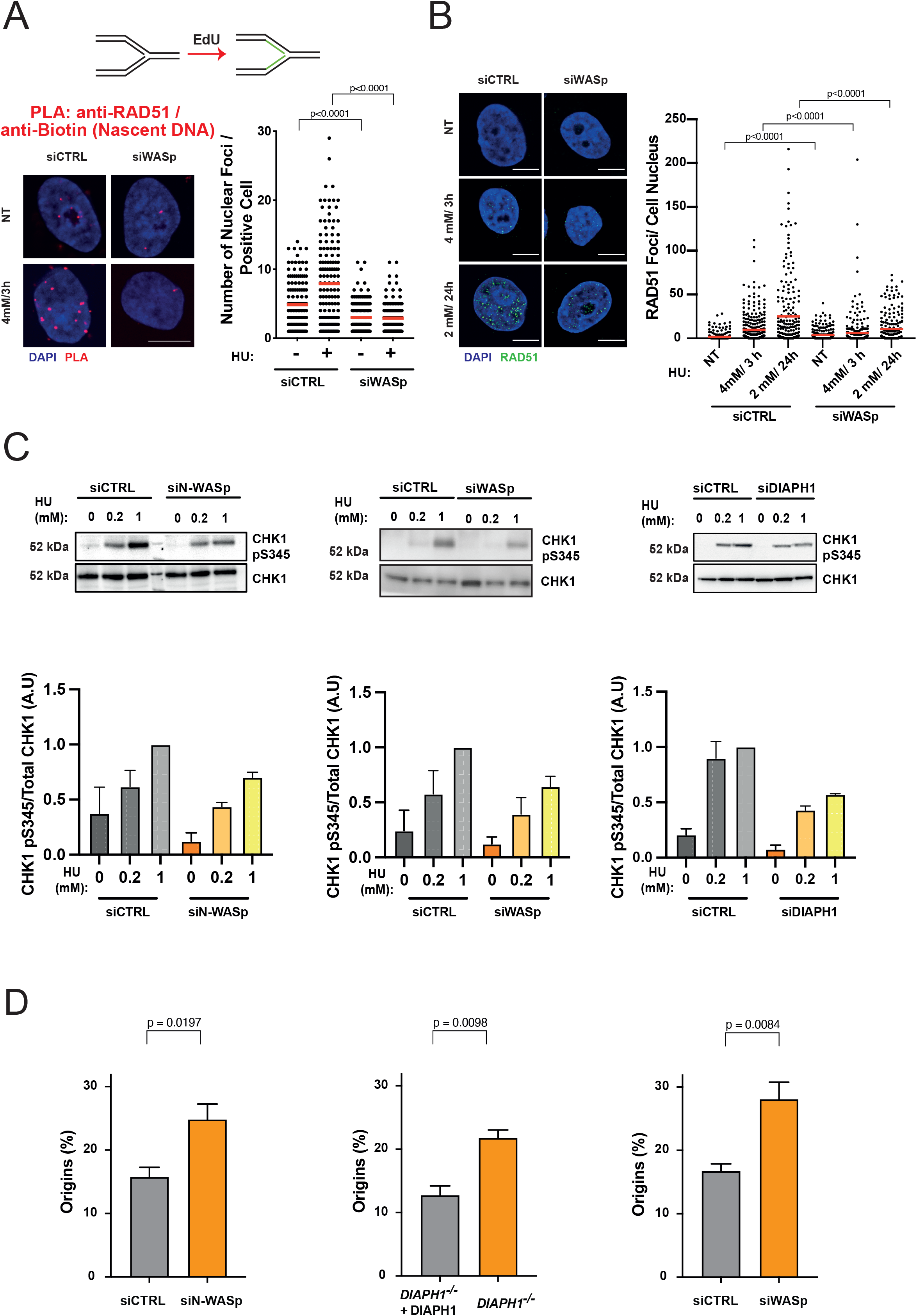
Depletion of WASp, N-WASp or DIAPH1 leads to defects in ATR signalling activation. A. Dot plots of number of RAD51/Biotin (Nascent DNA) PLA foci per nucleus in HeLa cells treated with control siRNA or siRNA against WASp upon treatment with 4 mM HU for 3 hours. Dot plots represent data pooled from two independent experiments (red lines indicate mean values). Statistical significance was determined using the Mann-Whitney test. B. Representative images and dot plots of the number of RAD51 foci per nucleus in HeLa cells treated with control siRNA or siRNA targeting WASp upon HU treatment. Dot plots represent data pooled from two independent experiments. Statistical significance was determined using the Mann-Whitney test. B. WB showing CHK1 phosphorylation in HeLa cells with or without indicated HU treatment for 1 h, treated with control siRNA (siCTRL) or siRNA targeting N-WASp or WASp as well as the same analysis in DIAPH1-deficient patient-derived cells and the corresponding control cells. Bar charts represent mean +/− SEM of three experiments and statistical significance was determined using t test. C. Quantification of the frequency of origin firing in HeLa cells treated with control siRNA or siRNA against N-WASp, WASp or DIAPH1. Bar charts represent mean +/− SEM of four (siN-WASp and siWASp) or three (DIAPH1) independent experiments. Statistical significance was determined using t test.

Since impaired formation of RPA foci and defective DNA replication was not restricted to WASp deficiency alone, but was observed with loss of multiple other members of the NPF/ANs protein family, we considered that this phenotype could arise through at least three distinct mechanisms: (i) an overall decrease in the levels of RPA expression in cells, (ii) defective nucleolytic processing of HU-perturbed forks failing to generate ssDNA, and/or (iii) defects in a downstream process that promotes RPA association with ssDNA. To distinguish between these possibilities, we first analysed total RPA2 protein levels in cells depleted of NPF/ANs, which indicated that loss of these factors does not affect RPA2 levels (Figure 6A), as we previously showed in WASp-deficient T cells^15^. We also used sub-cellular fraction and WB to confirm that the cytoplasmic and nuclear pools of RPA1 and 2 did not alter upon depletion of NPF/ANs (Figure S5A). Next, we analysed efficiency of ssDNA generation at HU-perturbed RFs by BrdU staining under non-denaturing condition. Here too, we did not observe a decrease in ssDNA generation (end-resection) at perturbed RFs upon depletion of these factors (Figure 6B). Thus, we considered it likely that NPF/ANs may function as RPA “chaperones” facilitating RPA loading and/or stability at RFs both, in *cis* as in case of WASp(15) but also in *trans*, likely via modulating the actin state, i.e., G-actin (monomeric) versus F-actin (polymeric).

**Figure 6.**
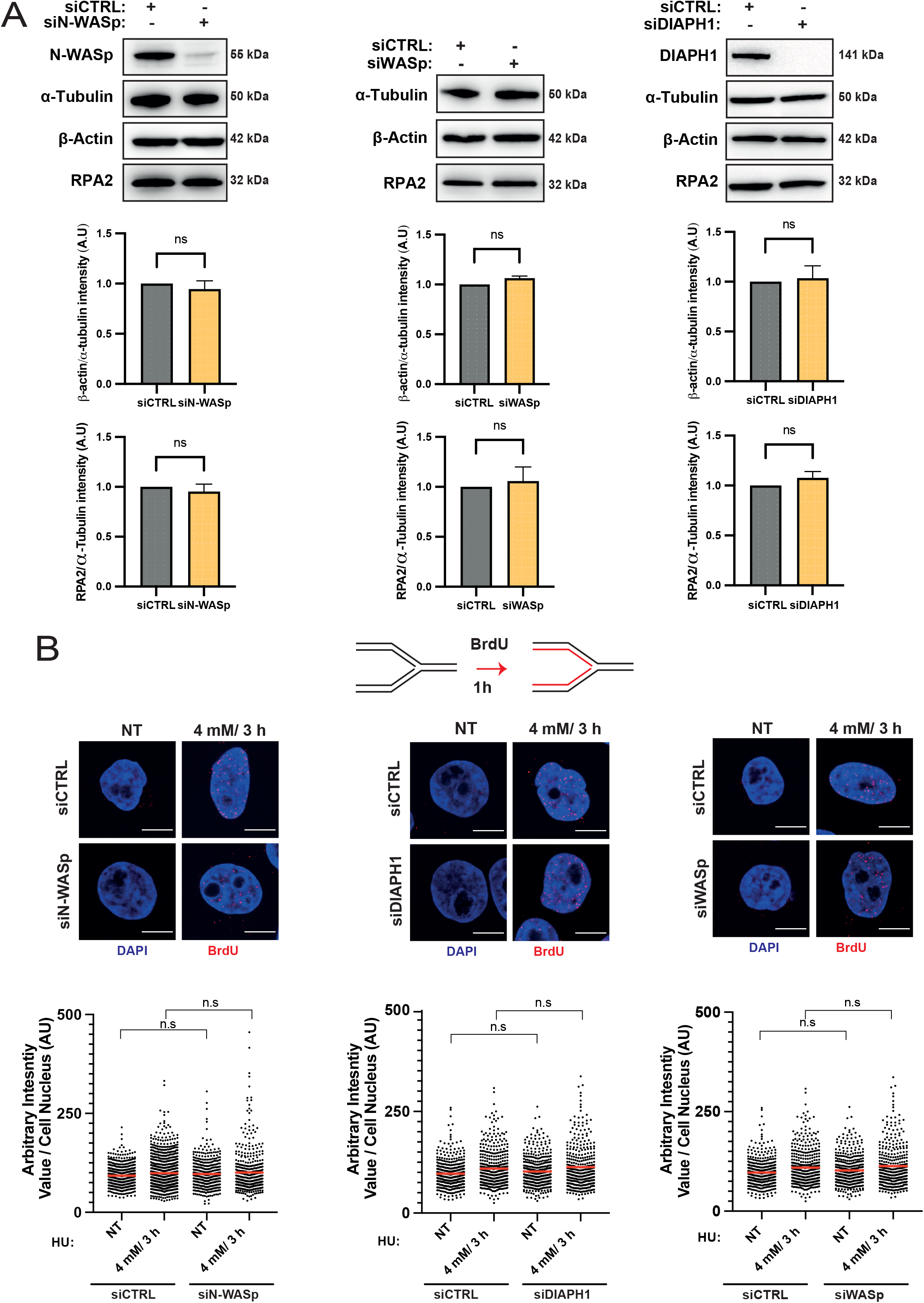
Depletion of WASp, N-WASp or DIAPH1 neither affects RPA or β-actin protein expression nor ssDNA formation. A. WB and quantitation showing protein levels of RPA2 and β–actin upon depletion of N-WASp, DIAPH1 or WASp. Bar charts represent mean +/− SEM of three experiments and statistical significance was determined using t test. B. Representative images and dot plots of BrdU signal intensity per nucleus of Hela cells treated with control siRNA (siCTRL) or siRNA targeting N-WASp, DIAPH1 or WASp upon HU treatment at the indicated dose/duration (red lines indicate mean values). Cells were incubated with BrdU (10 μM) for 1h prior to HU treatment. Dot plots represent data pooled from two independent experiments. Statistical significance was determined using the Mann-Whitney test. n.s., nonsignificant.

A simple prediction from this model would be that a surplus of RPA may be enough to overcome limitation associated with suboptimal RPA chaperoning in NPF/AN-deficient cells. To test this hypothesis, we utilised the previously characterised SuperRPA U2OS cells generated in the Lukas lab, that display a modest two-fold excess of all three RPA subunits. Importantly, these cells retain a normal cell-cycle profile and do not show spontaneous defects in either DNA replication or ATR activation(48). Strikingly, RPA over-expression largely restored RPA loading/stability at HU-perturbed forks in DIAPH1- or WASp-deficient cells (Supplementary Figure S5B), as well as restored global RF dynamics (Supplementary Figure S5C) with a concomitant rescue of ectopically-increased fork collapse – as measured by 53BP1 foci formation (Supplementary Figures S6A, B). To examine the specificity of the rescue phenotype we tested if a surplus of RPA can rescue, in general, any phenotype associated with RF instability. To this end, we analysed DNA replication in SuperRPA cells depleted for BRCA2, since BRCA2-depletion confers a strong defect in fork stability (44,49). In support of a specific role for NPF/ANs in facilitating RPA association with ssDNA and RF stability, we did not observe a similar rescue of BRCA2-associated replication phenotype in SuperRPA cells (Supplementary Figures S6C, D).

### Actin polymerisation mutants recapitulate the phenotypes associated with loss of NPF/ANs

Our above results suggest that actin nucleators, i.e., components of ARP2/3 or formin-dependent pathways, act early in the RSR pathway to promote the localisation of RPA to perturbed forks, stimulating efficient RPA-ssDNA complex formation both, directly through WASp-RPA interaction(15) but also indirectly through another putative mechanism. Indeed, a recent report by Pfitzer *et al*. (50) demonstrated that RPA2 coimmunoprecipitates (co-IPs) with nuclear actin, which proposes a model that G-actin/F-actin state itself may somehow influence the ability of RPA to form RPA-ssDNA complexes. To test this hypothesis, we first show that sepharose-coated monomeric G-actin directly binds purified RPA complex (Figure 7A and Supplementary Figure S7A) by *in-vitro* pull-down assays, suggesting that the *in vivo* RPA:G-actin association is also likely to be a direct interaction. Furthermore, we show that actin co-IPs with RPA2 in human cells spontaneously (without HU induced replication stress), however, upon HU treatment, this RPA:actin interaction is decreased (Supplementary Figure S7B). Notably, and in direct support of the role of NPF/ANs in “chaperoning” RPA-ssDNA complex formation, we observe that a hyper-depolymerising G13R β-actin mutant (YFP-NLS-β-actin^G13R^)(51,52) preferentially interacts with RPA2 in co-IP analyses from human HEK293TN cell extracts, when compared to WT (YFP-NLS-β-actin^WT^) or the S14C hyper-polymerising mutant of β-actin (YFP-NLS-β-actin^S14C^) (Figure 7B) (52). Accordingly, cells over-expressing an actin polymerization-incompetent (YFP-NLS-β-actin^G13R^) mutant that preferentially associates with RPA phenocopy the abnormal phenotypes associated with the loss of NPF/ANs i.e., defective RPA foci formation, ATR activation as well as global impairment in DNA replication (Figures 7C, D, Supplementary Figure S7C). In contrast, cells expressing actin polymerization-hypercompetent S14C mutant (YFP-NLS-β-actin^S14C^) showed phenotypes similar to cells expressing the control YFP-NLS-β-actin^WT^ (Figures 7C, D)

**Figure 7.**
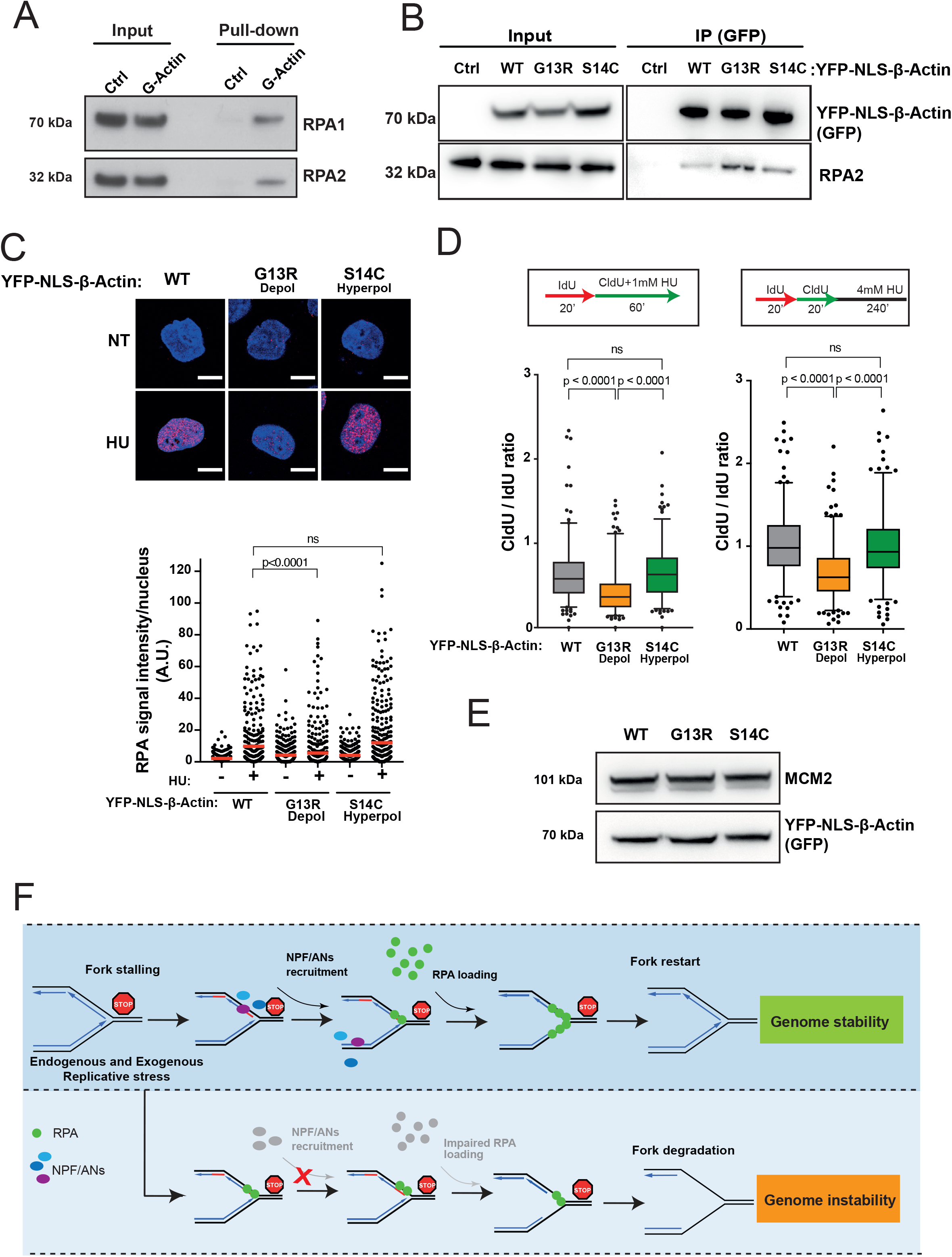
Monomeric actin directly binds RPA and actin polymerization is required for efficient RPA deposition at perturbed replication forks. A. RPA directly binds G-actin. WB of RPA *in vitro* pull down with G-actin conjugated sepharose beads. Ctrl, control. B. WB of immunoprecipitation (IP) experiments in HEK293TN cells expressing YFP-NLS-β-actin WT, G13R or S14C mutant. C. Representative images and dot plot of RPA signal intensity in HeLa cells expressing YFP-NLS-β-actin WT, G13R or S14C mutant upon 1mM HU treatment for 1h. Dot plots represent data pooled from three independent experiments. Statistical significance was determined using the Mann-Whitney test. ns, nonsignificant. D. Box plots of CldU/IdU tract ratios of HeLa cells expressing YFP-NLS-β-actin WT, G13R or S14C mutant proteins treated with 1mM HU during CldU labelling pulse (left panel) or CldU/IdU pulse labelling was followed by treatment with 4mM HU for 4h (right panel). Whiskers indicate 5-95 percentile. Box plots represent data pooled from two independent experiments. Statistical significance was determined using the Mann-Whitney test. ns, nonsignificant. E. Representative WB showing levels of expression of YFP-NLS-β-actin WT, G13R or S14C in HeLa cells from C and D. F. Proposed model for the role of NPF/ANs in the replicative stress response. Replicative stress leads to fork stalling and generation of ssDNA, subsequently the association of actin nucleators (ANs) and nucleation promoting factors (NPFs) with sites of ongoing replication facilitates RPA deposition, either directly (via WASp) or indirectly (via actin polymerization), in order to protect ssDNA generated at perturbed forks. This allows for an efficient fork restart and promotes genome stability. Depletion of NPF/ANs and/or interference with actin polymerization impairs RPA loading/stability on ssDNA leading to extensive nascent strand degradation and ultimately genome instability.

Note, WT and S14C mutant have a similar RPA foci phenotype (Figure 7C), suggesting a rate-limiting step within the actin pathway that is distinct from actin filament formation, especially since we also show that all 3 actins (WT, S14C, G13R) are expressed at a similar level (Figure 7E). Thus, our data suggest a unique functional and cooperative interaction between RPA, actin and NPF/ANs where a combination of *cis* and *trans*, actin-state dependent, mechanisms calibrate RPA activity at the forks to safeguard DNA replication (Model, Figure 7F).

## DISCUSSION

The RPA-ssDNA-protein complex plays a key role in protecting ssDNA generated during DNA metabolism, including at RFs, as well as acting as a “platform” for recruitment/signalling/regulation of a plethora of DNA repair factors. However, increased RPA levels can be toxic likely due to excessive formation of RPA-ssDNA complexes (14,48,53–55). Our recent report(15) as well as work in yeast *Saccharomyces cerevisiae* (Sc) and Xenopus together point to active mechanisms employed by cells to dynamically control the ability of RPA to associate/dissociate with ssDNA (10,11).

Here we have shown that not only WASp, but also other members of the NPF/ANs protein family such as N-WASp, ARP2/3, or DIAPH1 as well as actin itself, promote efficient localisation of RPA to perturbed forks and by doing so promote ATR activation and supress ectopic fork resection. In line with this, we show that NPF/ANs involved in ARP2/3 or formin-dependent actin polymerisation are specifically enriched at perturbed forks and their loss preclude RPA localisation to ssDNA resulting in defective activation of ATR, excessive fork degradation and ultimately RF collapse. Furthermore, factors from the two actin polymerisation pathways (branched and unbranched) act in parallel, as combined disruption of both pathways results in an increased inability of cells to protect RFs. Accordingly, our work reveals an unexpected yet highly significant association between actin nucleation and the ability of cells to respond to replicative stress. Notably, our data indicate that actin interacts directly with RPA and overexpression of a nuclear localizing actin mutant that impedes its polymerisation (G13R)(56) leads to defective RPA foci formation and replication fork instability but the polymerization-hypercompetent actin mutant (S14C) does not(51,57). Our findings with G-actin (G13R) and F-actin (S14C) mutants on RPA foci formation, ATR activation, and efficiency of DNA replication (Figure 7 and Supplementary Figure S7C) suggest that the role of NPF/ANs in this process may be at least by two complementary mechanisms: (i) in *trans*, via control of the actin polymerisation and (ii) in *cis*, via WASp-RPA binding(15). Since β-actin is a sub-unit of multiple chromatin-remodelling complexes,(58–60) and their disruption causes global chromatin reorganization, this proposes an additional mechanism that may potentially contribute to the dysfunctional replication stress response (RSR) in cells lacking NPF/ANs.

Accordingly, our findings propose a model in which localized actin assembly at replication forks provides a spatiotemporal coordination of the RSR by delivering a pool of RPA at the right time (replicative stress) and at the right place (perturbed RFs). This could perhaps involve a “hand-off” like mechanism to transfer RPA from low-affinity sites on polymerized actin (F-actin) to high-affinity sites on ssDNA. Equally, it is possible that monomeric actin (G-actin) serves as an RPA rheostat, limiting the pool of free RPA to suppress toxicity associated with supra-physiological loading of RPA onto ssDNA. Upon fork stalling (RSR signaling) change in actin state (actin polymerization) may drive the dissociation of G-actin-bound RPA, which increases the pool of RPA available for binding to ssDNA and facilitates, both *in cis* and *in trans*, the efficient formation of RPA-ssDNA complexes, essential for fork protection and robust checkpoint activation (15). As such, a role for ATR in promoting nuclear actin filament formation has been established (21). In this context, our findings raise an intriguing possibility that an ATR-dependent feed-forward loop links the control of RPA assembly with the severity of replicative stress. Such a mechanism would allow for “discreet” activation of the replication checkpoint, facilitating “local”*vs* “global” responses. Indeed, the key function of checkpoint signaling is to control the global surplus of nuclear RPA, shielding RFs from catastrophic breakage. (61) Our findings insert NPF/ANs into this fundamental process in both immune and non-immune cell types.

Patient phenotypes with mutations in actin polymerisation pathways closely resemble those found in genetic disorders with defects in replication/DNA repair pathways i.e., increased cancer risk, failure of the hematopoietic system (immunodeficiency), microcephaly, neurodegeneration, developmental delay, or premature aging (16,62–65). The severity and broad spectrum of these phenotypes likely reflect context- and cell-dependent variability, including the high redundancy within these complimentary mechanisms, as well as potential functions of NPF/ANs outside of the RSR/DDR pathways. Indeed, disruption of WASp:ARP2/3- and DIAPH1-dependent actin polymerisation pathways confers a stronger replication defect than loss of a single pathway.

Altogether, our findings shed new light on the molecular mechanism by which actin nucleators and nucleation-promoting factors facilitate efficient DNA replication and suppress genome instability during the RSR, and further clarify the pathophysiology of human disorders linked to WASp and DIAPH1 deficiencies.

## Data Availability

Unprocessed imaging files generated for this study and source data for STRING analysis of iPOND have been deposited on Mendeley Database (DOI: 10.17632/gvcgdt5dc5.1).

## Acknowledgements

We thank L. I. Toledo (University of Copenhagen) for U2OS SuperRPA cell line, M. Seidman and M. Bellini (NIH, Baltimore, USA) for scholarly input, M. Dylewska for technical assistance, and the Flow Cytometry & Light Microscopy Facility at the ICR (K. Betteridge, A. Stojiljkovic, H. Ale and M. Guelbert) for assistance with microscopy and flow cytometry cell sorting.

## Contributions

W.N. and Y.M.V. conceived the project. W.N., J.N. and P.M. designed and analysed the experiments with contribution from R.B. J.N., P.M. and R.B. performed most of the experiments. J.K., A.K., C.M, C.S, V.B and K.K. contributed with specific experiments. A.A. provided advice on project design and data analysis. C.B. provided assistance with visualisation of actin polymers and high content imaging. G.S.S. provided advice on project design and carried out initial analysis and complementation of DIAPH1 cell line with assistance from B.L.W.; F.S. provided the DIAPH1 cell line. W.N. wrote the manuscript with inputs from Y. M.V., and editing contributions from J.N., P.M., R.B., A.A and G.S.S., and all authors reviewed it.

The authors declare no competing interests

