## Supplementary figures for "Actin nucleators safeguard replication forks by limiting nascent strand degradation"

**Figure S1**

A. Schematic showing iPOND experimental design.

B. WB analysis of iPOND in HEK293TN cells showing DIAPH1 and N-WASp association with nascent DNA (EdU) upon exposure to HU with/without thymidine (Thy) chase. PCNA is shown as a reference event.

C. WB analysis of iPOND in human T cells showing DIAPH1 and WASp association with nascent DNA (EdU) upon exposure to HU with/without thymidine (Thy) chase. upon exposure to HU. RPA2 is shown as a reference event.

D. Representatives images of antibody controls for PLA experiments; anti Biotin only (rabbit or mouse, as indicated) and anti-PCNA only act as negative controls, anti-Biotin (rabbit):anti-Biotin (mouse) act as a positive control.

E. Representative images and dot plots of number of N-WASp/PCNA PLA and WASp/PCNA PLA foci per nucleus in HeLa cells untreated (NT) or treated with HU as indicated (red lines indicate mean values). Dot plots represent data pooled from two independent experiments. Statistical significance was determined using the Mann-Whitney test.

**Figure S2**

A. WB analysis confirming DIAPH1 and N-WASp knock-down efficiency in HeLa cells.

B. Bar chart representing qPCR data confirming WASp knock-down efficiency in HeLa cells.

C. qPCR and WB analysis confirming WASp and N-WASp knock down efficiency in HeLa cells with additional siRNAs (denoted as #1 or #2) as in panel D.

D. Box plots of CldU/IdU tract ratios from HeLa cells treated with control siRNA or siRNA targeting WASp or N-WASp as indicated. Cells were treated with 1mM HU during CldU labelling pulse. Whiskers indicate 5-95 percentile. Box plots represent data pooled from two independent experiments. Statistical significance was determined using the Mann-Whitney test.
E. Box plots of CldU/IdU tract ratios from HeLa cells treated with CK-666 (ARP2/3 inhibitor) (+) or DMSO (-). Cells were treated with 1mM HU during CldU labelling pulse. Whiskers indicate 5-95 percentile. Box plots represent data pooled from three independent experiments. Statistical significance was determined using the Mann-Whitney test.

**Figure S3**

A. Box plots of CldU/IdU tract ratios from HeLa cells depleted of WASp (siWASp) or not (siCTRL), and treated with CK-666 ARP2/3 inhibitor (+) or DMSO (-). Cells were exposed to 1mM HU during CldU labelling pulse. Whiskers indicate 5-95 percentile. Box plots represent data pooled from three independent experiments. Statistical significance was determined using the Mann-Whitney test.
B. Representative images and dot plots of the number of 53BP1 foci per cyclin A positive HeLa cells, treated with CK-666 (ARP2/3 inhibitor) or CTRL (DMSO), and DIAPH1-deficient patient-derived cells exposed to HU (+) or not (-). Dot plots represent data pooled from two independent experiments. Statistical significance was determined using the Mann-Whitney test.
C. Representative images and quantification of micronuclei formation (indicated by an arrow) in HeLa cells treated with CK-666 (ARP2/3 inhibitor) (+) or with DMSO (-).

Bar charts represent mean +/- SEM of three independent experiments. Statistical significance was determined using t test.

D. Box plots of CldU/IdU tract ratios from HeLa cells treated with control siRNA or siRNA targeting WASp mock treated or treated with Mirin or siRNA targeting DNA2. CldU/IdU pulse labelling was followed by treatment with 4mM HU for 4h. Whiskers indicate 5-95 percentile. Box plots represent data pooled from two independent experiments. Statistical significance was determined using the Mann-Whitney test.

**Figure S4**

A. Representative images and dot plots of RPA signal intensity per nucleus of DIAPH1-deficient patient-derived cells exposed to HU at the indicated dose/duration (red lines indicate mean values). Dot plots represent data pooled from two independent experiments. Statistical significance was determined using the Mann-Whitney test. A.U., arbitrary units.
B. Representative images and dot plots of RPA signal intensity per nucleus of Hela cells treated with control siRNA or additional siRNA targeting WASp or N-WASP upon HU treatment as indicated (red lines indicate mean values). Dot plots represent data pooled from two independent experiments. Statistical significance was determined using the Mann-Whitney test.
C. Representative images and dot plots of RPA signal intensity per nucleus of HeLa cells treated with CK-666 (ARP2/3 inhibitor) (+) or DMSO (-) upon HU treatment as indicated (red lines indicate mean values). Dot plots represent data pooled from two independent experiments. Statistical significance was determined using the Mann-Whitney test.

**Figure S5**

A. WB showing cytoplasmic (α-tubulin as cytoplasmic loading control) and nuclear protein levels (Lamin-B1 as nuclear loading control) of RPA1 and RPA2 upon depletion of N-WASp, WASp or DIAPH1.

B. Representative images and dot plots of RPA signal intensity per nucleus of U2OS WT cells and U2OS cells overexpressing RPA (RPA O.E) treated with control siRNA (CTRL) or siRNA targeting DIAPH1 or WASp upon HU treatment or no treatment (NT) as indicated (red lines indicate mean values). Dot plots represent data pooled from three or two independent experiments for DIAPH1 and WASp, respectively. Statistical significance was determined using the Mann-Whitney test. ns, nonsignificant.

C. Box plots of CldU/IdU tract ratios of U2OS WT cells and U2OS cells overexpressing RPA (RPA O.E) treated with control siRNA (-) or siRNA targeting WASp or DIAPH1 (+) and exposed to 1mM HU during CldU labelling pulse. Whiskers indicate 5-95 percentile. Box plots represent data pooled from two independent experiments. Statistical significance was determined using the Mann-Whitney test.

**Figure S6**

A. Representative images and dot plots of the number of 53BP1 foci per cell nucleus in U2OS WT cells and U2OS cells overexpressing RPA upon depleting DIAPH1 or WASp (siDIAPH1 or siWASp) or their control knock down (siCTRL) (red lines indicate mean values). Dot plots represent data pooled from two independent experiments for DIAPH1 and WASp. Statistical significance was determined using the Mann-Whitney test.

B. qPCR and WB analysis confirming WASp and DIAPH1 knock-down efficiency in U2OS WT cells and U2OS cells overexpressing RPA.
C. Representative images and dot plots of RPA signal intensity per nucleus of U2OS WT cells and U2OS cells overexpressing RPA treated with control siRNA or siRNA targeting BRCA2 upon HU treatment as indicated (red lines indicate mean values). Dot plots represent data pooled from two independent experiments.
D. Box plot of CldU/IdU tract ratios from U2OS WT cells and U2OS cells overexpressing RPA treated with control siRNA or siRNA targeting BRCA2. CldU/IdU pulse labelling was followed by treatment with 4mM HU for 4 h. Whiskers indicate 5-95 percentile. Box plots represent data pooled from two independent experiments. Statistical significance was determined using the Mann-Whitney test.

**Figure S7**

A. Coomassie stained SDS-PAGE gel showing purified RPA complex and molecular weight (MW) marker.
B. WB of immunoprecipitation experiments in HEK293TN cells expressing YFP-NLS-β-actin WT treated or not with 4mM HU / 3 h.
C. WB showing CHK1 phosphorylation upon indicated HU treatment for 1 h in HeLa cells expressing YFP- NLS-β-actin WT and G13R mutant.

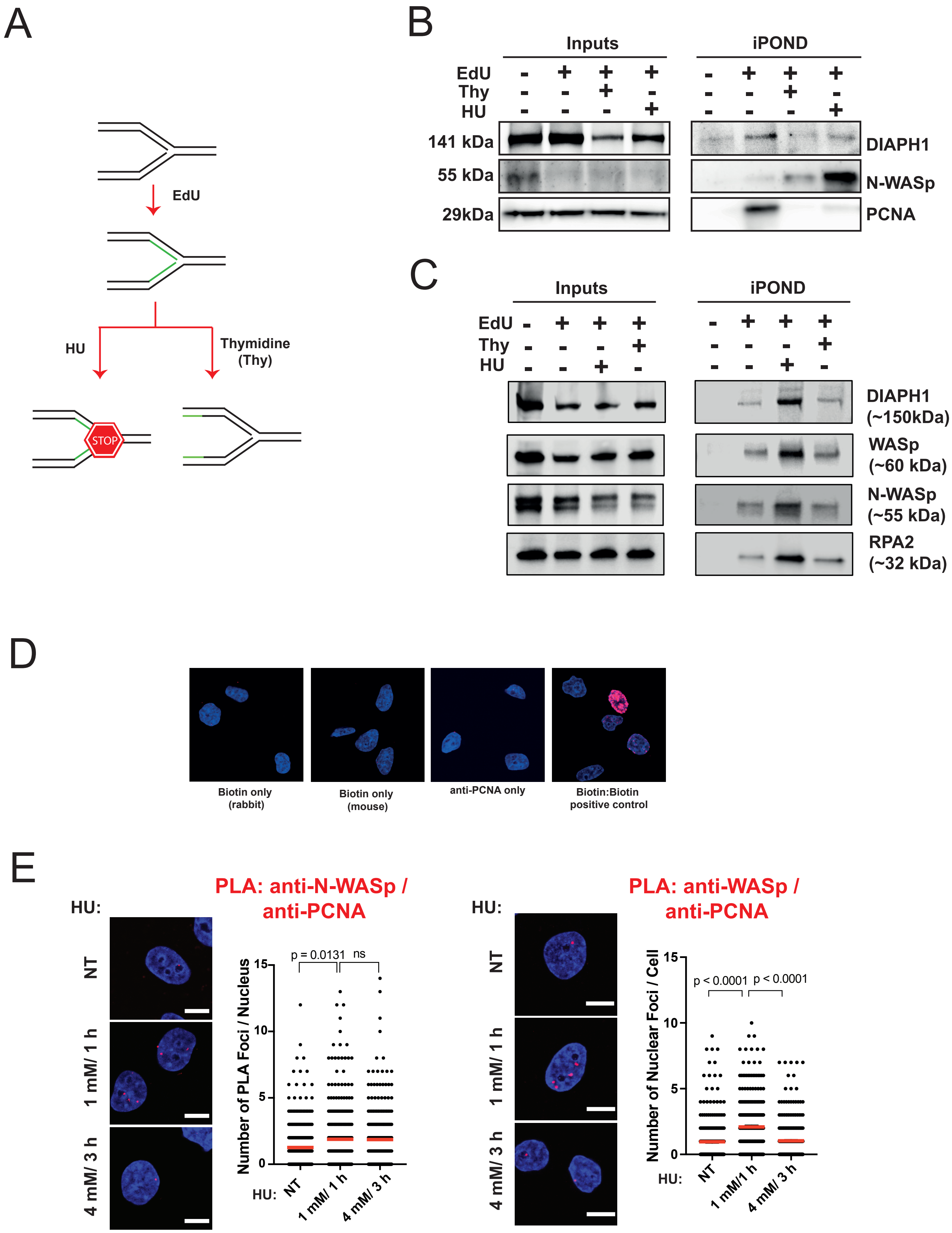

A
EdU
HU
Thymidine (Thy)
STOP
B
Inputs
iPOND
EdU
Thy
HU
141 kDa
55 kDa
29kDa
DIAPH1
N-WASp
PCNA
C
Inputs
iPOND
EdU
Thy
HU
DIAPH1 (~150kDa)
WASp (~60 kDa)
N-WASp (~55 kDa)
RPA2 (~32 kDa)
D
Biotin only (rabbit)
Biotin only (mouse)
anti-PCNA only
Biotin:Biotin positive control
E
PLA: anti-N-WASp / anti-PCNA
HU:
NT
1 mM/ 1 h
4 mM/ 3 h
p = 0.0131
ns
Number of PLA Foci / Nucleus
PLA: anti-WASp / anti-PCNA
p < 0.0001
p < 0.0001
Number of Nuclear Foci / Cell
1 mM/1 h
4 mM/ 3 h


Figure S1

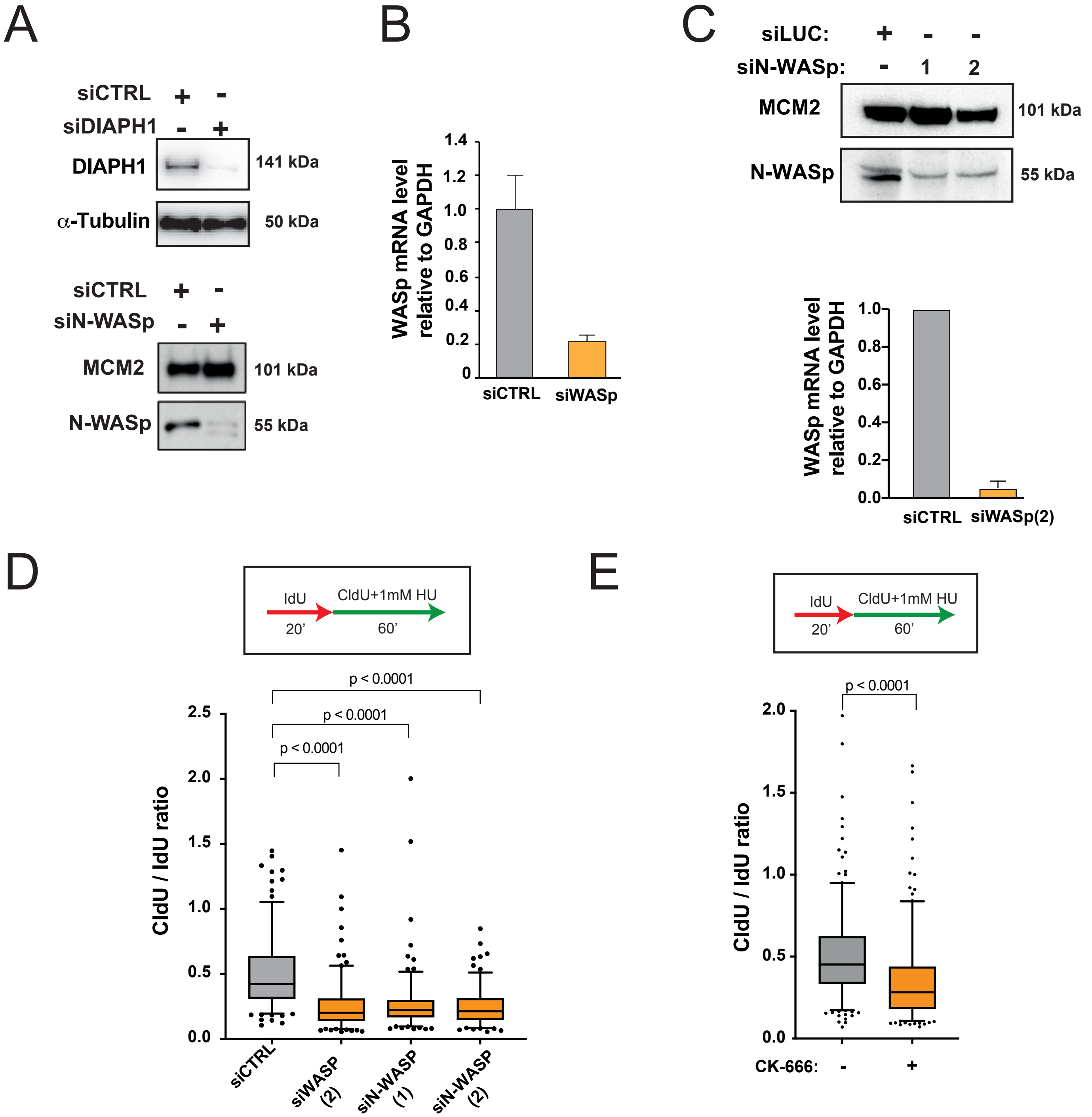
A
siCTRL + -
siDIAPH1 - +
DIAPH1
141 kDa
α-Tubulin
50 kDa
siCTRL + -
siN-WASp - +
MCM2
101 kDa
N-WASp
55 kDa
B
WASp mRNA level relative to GAPDH
1.4
1.2
1.0
0.8
0.6
0.4
0.2
0
siCTRL
siWASp
C
siLUC: + - -
siN-WASp: - 1 2
MCM2
101 kDa
N-WASp
55 kDa
WASp mRNA level relative to GAPDH
1.0
0.8
0.6
0.4
0.2
0.0
siCTRL
siWASp(2)
D
IdU
CldU+1mM HU
20'
60'
p < 0.0001
p < 0.0001
p < 0.0001
CldU / IdU ratio
2.5
2.0
1.5
1.0
0.5
0.0
siCTRL
siWASP (2)
siN-WASP (1)
siN-WASP (2)
E
IdU
CldU+1mM HU
20'
60'
p < 0.0001
CldU / IdU ratio
2.0
1.5
1.0
0.5
0.0
CK-666: - +

Figure S2

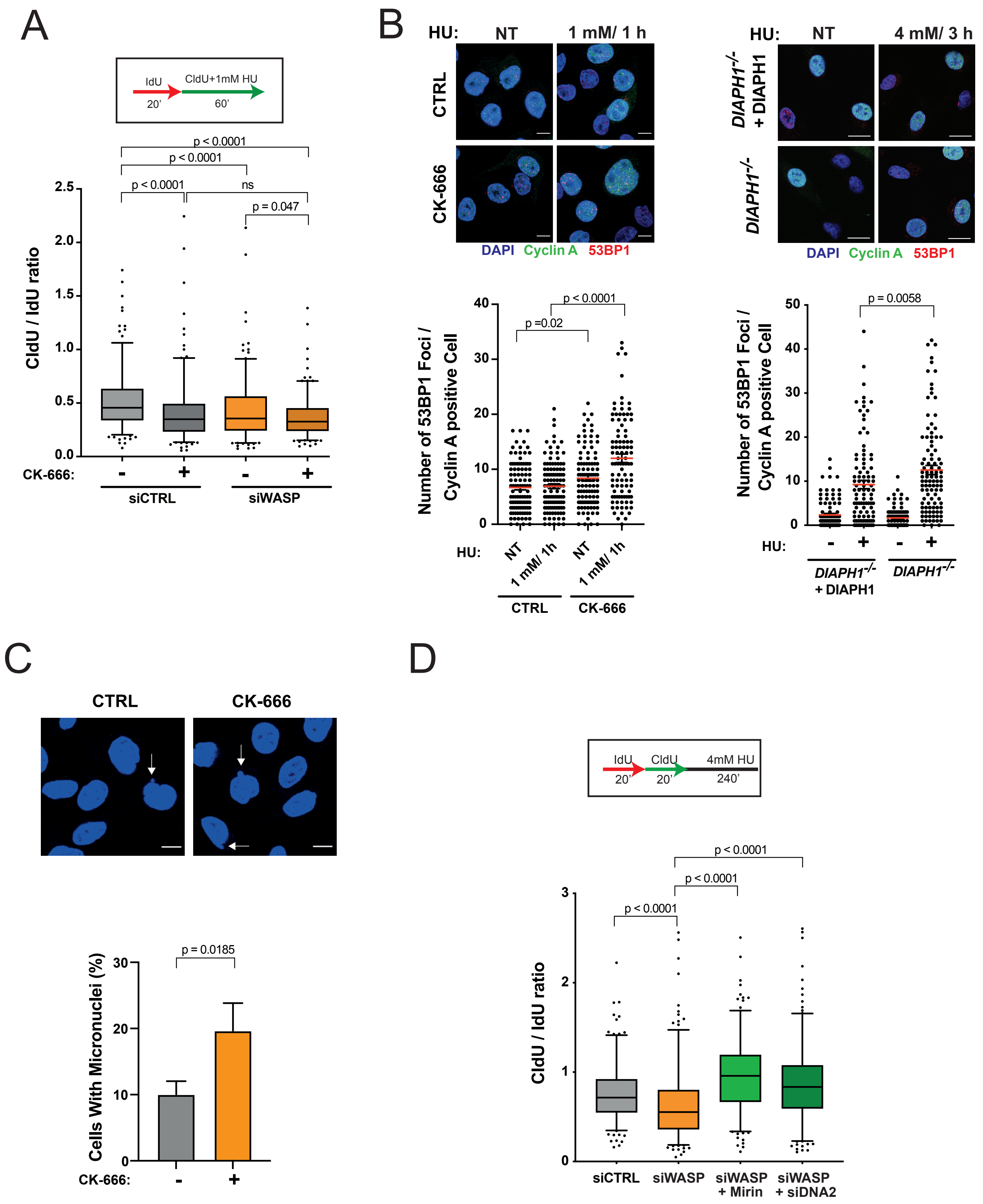

A
IdU
20'
CldU+1mM HU
60'
p < 0.0001
p < 0.0001
p < 0.0001
ns
p = 0.047
CldU / IdU ratio
CK-666:
siCTRL
siWASP
B
HU:
NT
1 mM/ 1 h
CTRL
CK-666
DAPI Cyclin A 53BP1
4 mM/ 3 h
DIAPH1-/- + DIAPH1
DIAPH1-/-
Number of 53BP1 Foci / Cyclin A positive Cell
p =0.02
p < 0.0001
p = 0.0058
1 mM/ 1h
C
CTRL
CK-666
Cells With Micronuclei (%)
p = 0.0185
D
IdU
20'
CldU
20'
4mM HU
240'
siCTRL
siWASP
siWASP + Mirin
siWASP + siDNA2


Figure S3

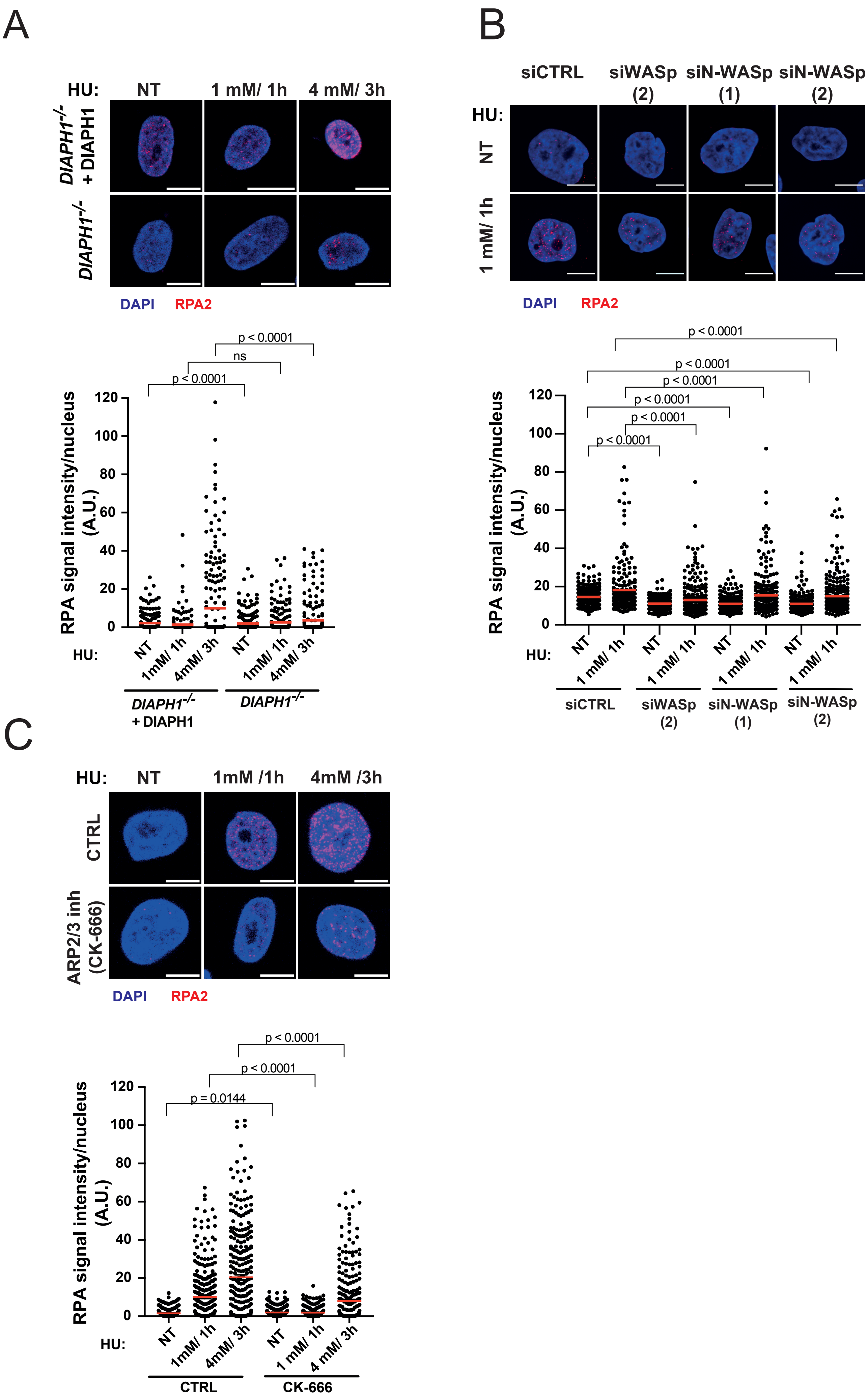
A
HU: NT 1 mM/ 1h 4 mM/ 3h
DIAPH1-/- + DIAPH1
DIAPH1-/-
DAPI RPA2
p < 0.0001
ns
p < 0.0001
RPA signal intensity/nucleus (A.U.)
120
100
80
60
40
20
0
HU: NT 1mM/ 1h 4mM/ 3h NT 1mM/ 1h 4mM/ 3h
DIAPH1-/- + DIAPH1
DIAPH1-/-
B
siCTRL siWASp (2) siN-WASp (1) siN-WASp (2)
HU: NT
1 mM/ 1h
DAPI RPA2
p < 0.0001
p < 0.0001
p < 0.0001
p < 0.0001
p < 0.0001
p < 0.0001
RPA signal intensity/nucleus (A.U.)
120
100
80
60
40
20
0
HU: NT 1 mM/ 1h NT 1 mM/ 1h NT 1 mM/ 1h NT 1 mM/ 1h
siCTRL siWASp (2) siN-WASp (1) siN-WASp (2)
C
HU: NT 1mM /1h 4mM /3h
CTRL
ARP2/3 inh (CK-666)
DAPI RPA2
p < 0.0001
p < 0.0001
p = 0.0144
RPA signal intensity/nucleus (A.U.)
120
100
80
60
40
20
0
HU: NT 1mM/ 1h 4mM/ 3h NT 1 mM/ 1h 4 mM/ 3h
CTRL
CK-666

Figure S4

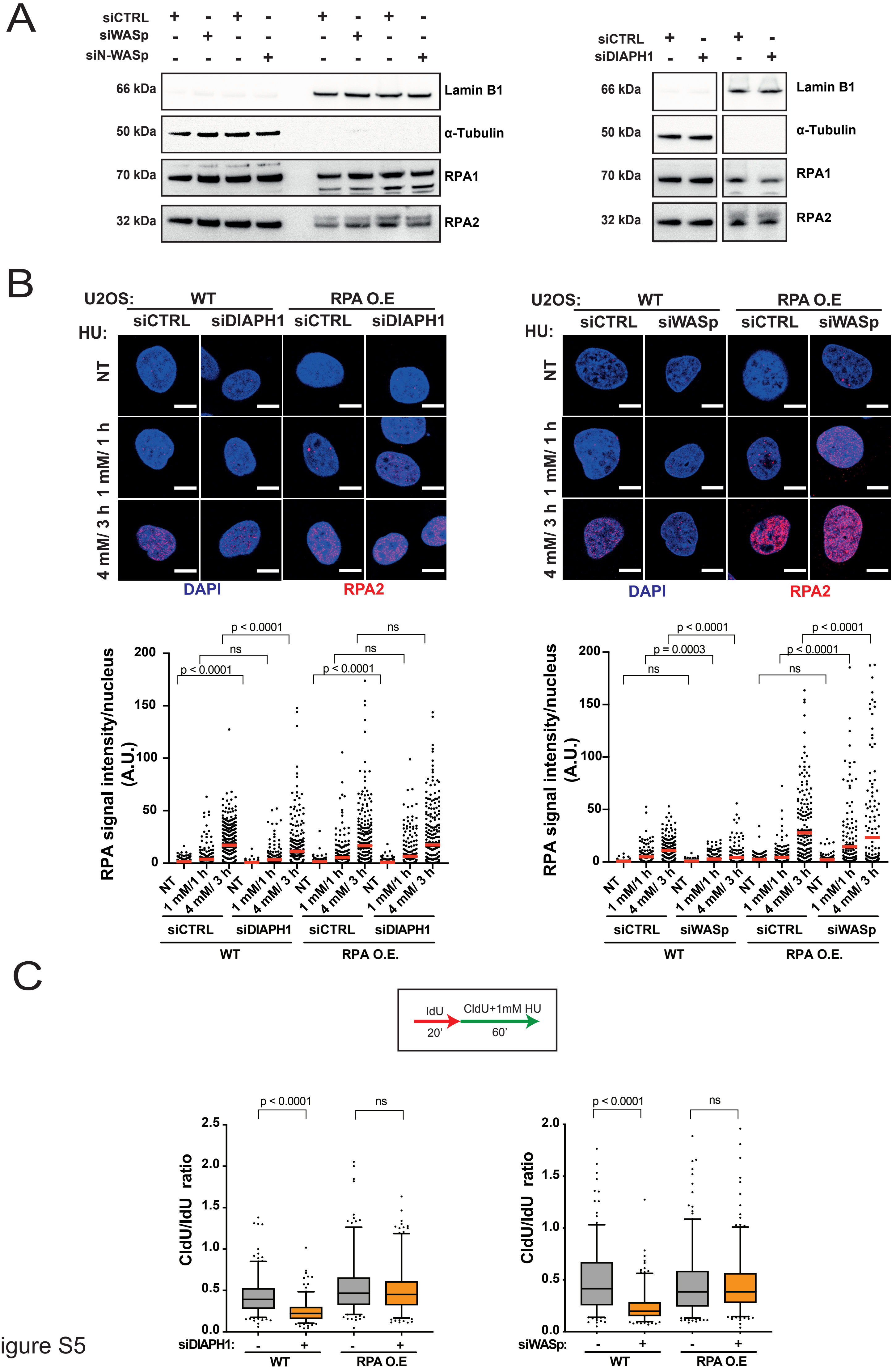

A
siCTRL
siWASp
siN-WASp
66 kDa
50 kDa
70 kDa
32 kDa
Lamin B1
α-Tubulin
RPA1
RPA2
siDIAPH1
B
U2OS:
WT
RPA O.E
HU:
NT
1 mM/ 1 h
4 mM/ 3 h
DAPI
RPA2
RPA signal intensity/nucleus (A.U.)
p < 0.0001
ns
p = 0.0003
RPA O.E.
C
IdU
20'
CldU+1mM HU
60'
CldU/IdU ratio
siDIAPH1:
siWASp:

Figure S5

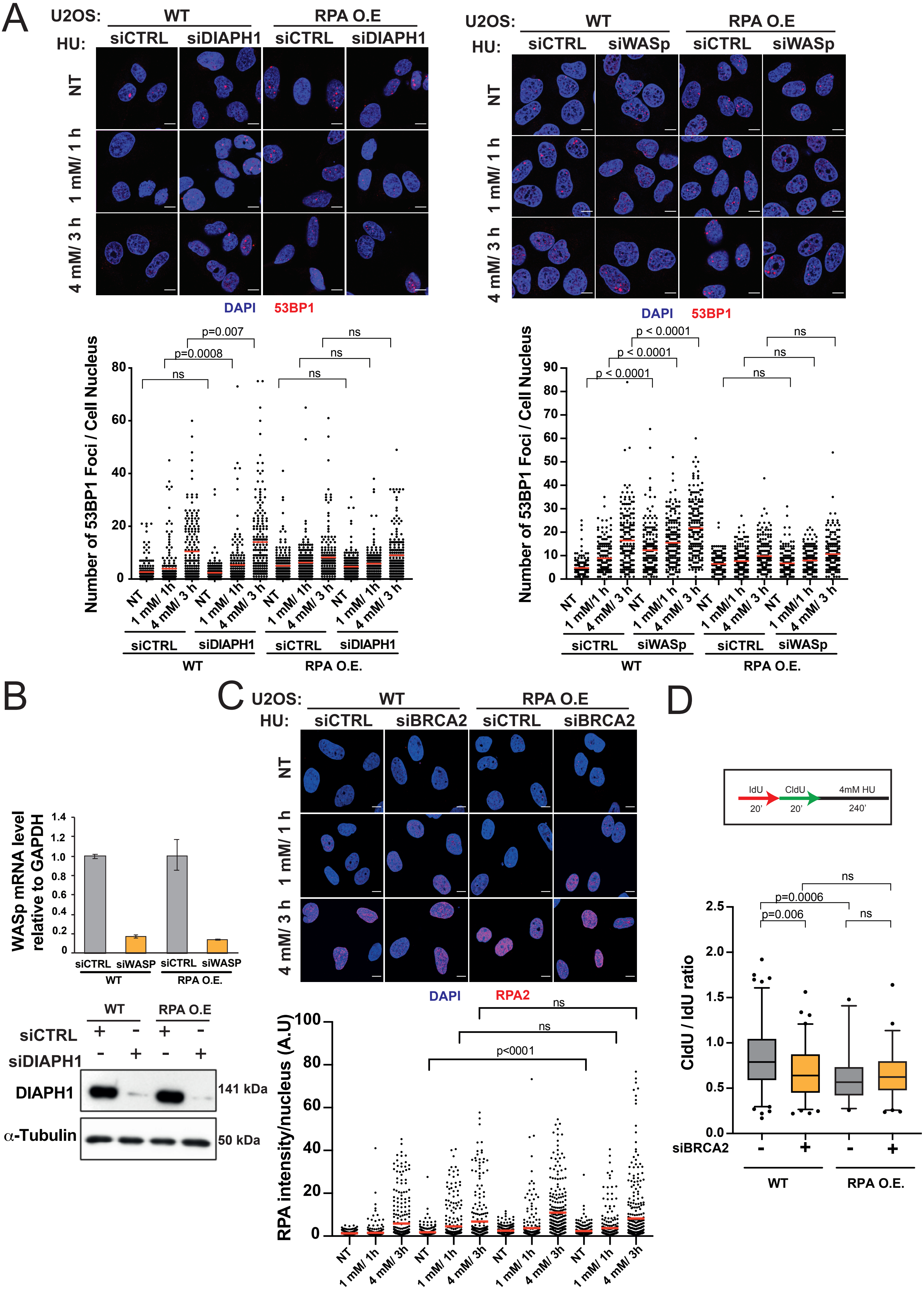
A
U2OS: WT RPA O.E
HU: siCTRL siDIAPH1 siCTRL siDIAPH1
NT
1 mM/ 1 h
4 mM/ 3 h
DAPI 53BP1
U2OS: WT RPA O.E
HU: siCTRL siWASp siCTRL siWASp
DAPI 53BP1
Number of 53BP1 Foci / Cell Nucleus
p=0.007
p=0.0008
ns
p < 0.0001
WT
RPA O.E.
B
WASp mRNA level relative to GAPDH
siCTRL siWASP siCTRL siWASP
siCTRL
siDIAPH1
DIAPH1
141 kDa
α-Tubulin
50 kDa
C
U2OS: WT RPA O.E
HU: siCTRL siBRCA2 siCTRL siBRCA2
DAPI RPA2
RPA intensity/nucleus (A.U)
p<0001
ns
D
IdU CldU 4mM HU
20' 20' 240'
CldU / IdU ratio
p=0.006
p=0.0006
ns
siBRCA2
WT
RPA O.E.

Figure S6

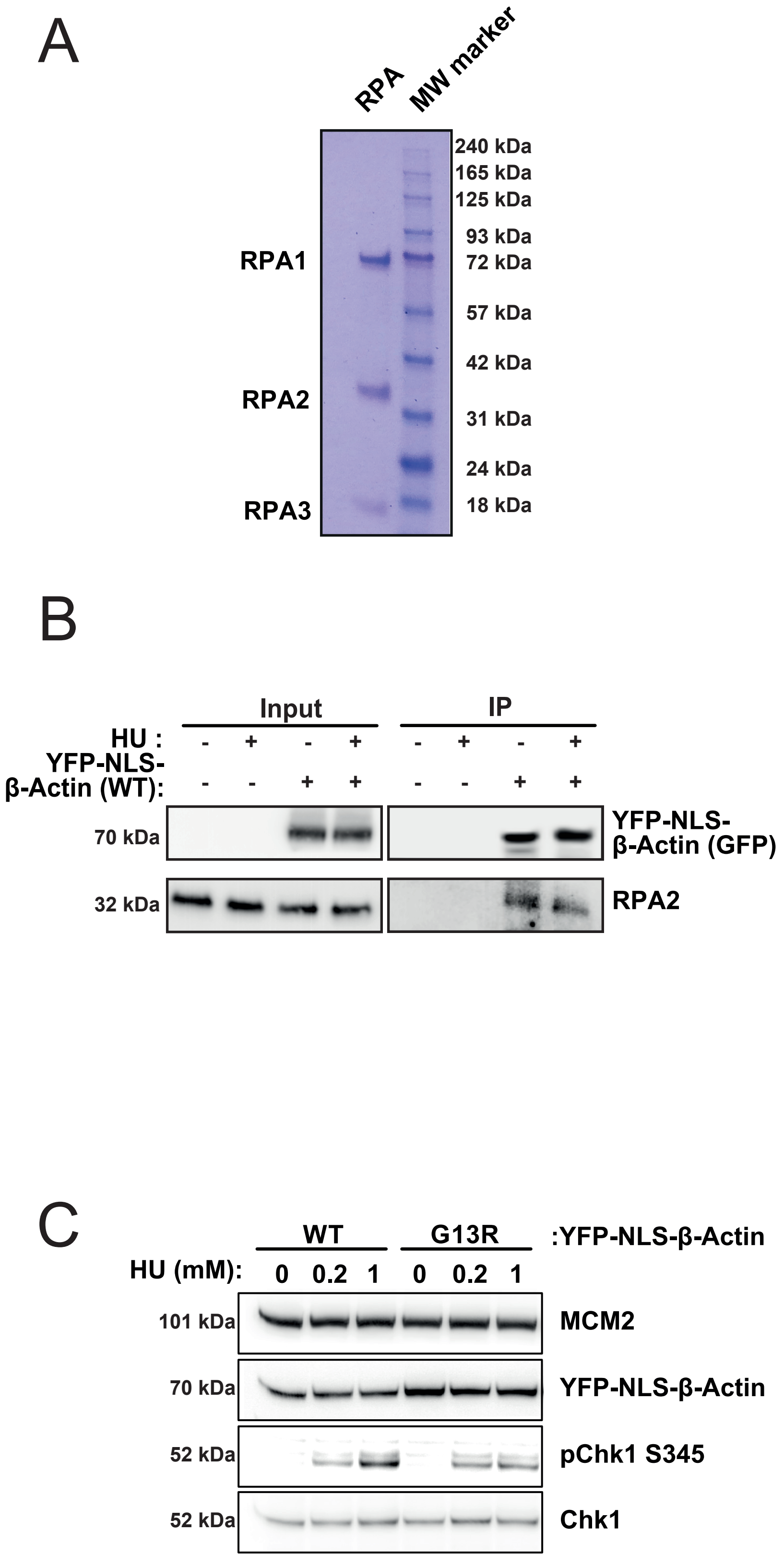
A
RPA
MW marker
240 kDa
165 kDa
125 kDa
93 kDa
72 kDa
57 kDa
42 kDa
31 kDa
24 kDa
18 kDa
RPA1
RPA2
RPA3
B
Input
IP
HU :
YFP-NLS-β-Actin (WT):
70 kDa
32 kDa
YFP-NLS-β-Actin (GFP)
RPA2
C
WT
G13R
:YFP-NLS-β-Actin
HU (mM):
0 0.2 1 0 0.2 1
101 kDa
70 kDa
52 kDa
52 kDa
MCM2
YFP-NLS-β-Actin
pChk1 S345
Chk1

Figure S7
